# Improved Protein Model in SPICA Force Field

**DOI:** 10.1101/2023.09.15.557852

**Authors:** Teppei Yamada, Yusuke Miyazaki, Shogo Harada, Ashutosh Kumar, Stefano Vanni, Wataru Shinoda

## Abstract

The previous version of the SPICA coarse-grained (CG) force field (FF) protein model focused primarily on membrane proteins and successfully reproduced the dimerization free energies of several transmembrane helices and stable structures of various membrane protein assemblies. However, that model had limited accuracy when applied to other proteins, such as intrinsically disordered proteins (IDPs) and peripheral proteins, because the dimensions of the IDPs in an aqueous solution were too compact, and protein binding on the lipid membrane surface was over-stabilized. To improve the accuracy of the SPICA FF model for the simulation of such systems, in this study we introduce protein secondary structure-dependent nonbonded interaction parameters to the backbone segments and re-optimize almost all nonbonded parameters for amino acids. The improved FF proposed here successfully reproduces the radius of gyration of various IDPs, the binding sensitivity of several peripheral membrane proteins, and the dimerization free energies of several transmembrane helices. The new model also shows improved agreement with experiments on the free energy of peptide association in water. In addition, an extensive library of nonbonded interactions between proteins and lipids, including various glycerophospholipids, sphingolipids, and cholesterol, allows the study of specific interactions between lipids and peripheral and transmembrane proteins. Hence, the new SPICA FF (version 2) proposed herein is applicable with high accuracy for simulating a wide range of protein systems.

**Graphical abstract:** 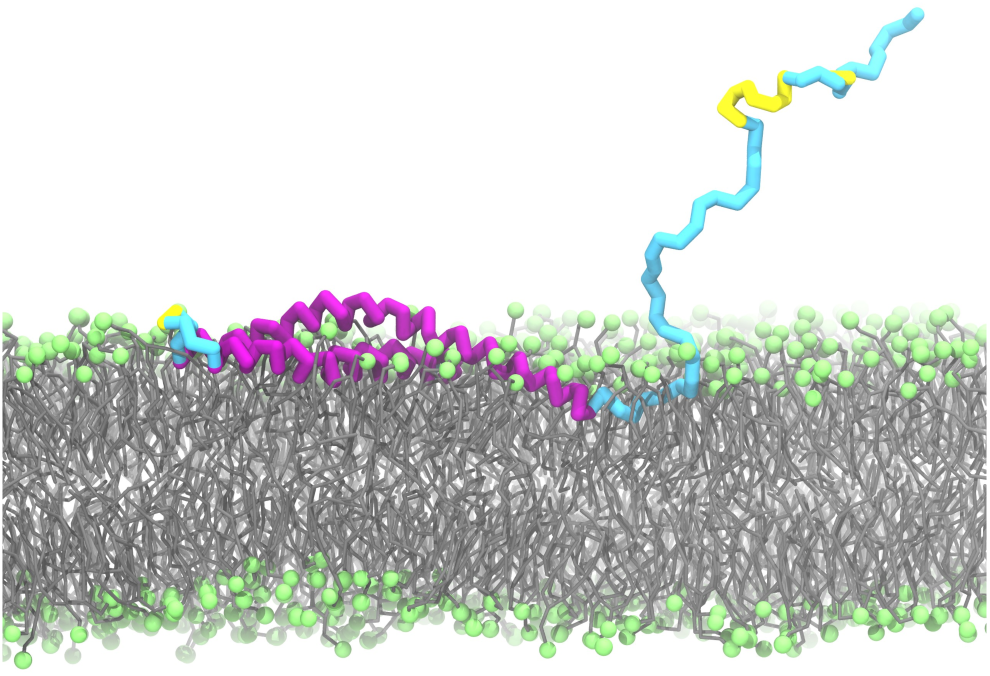

## 1. INTRODUCTION

Molecular dynamics (MD) simulations are crucial for studying the structural and dynamic details of proteins at high spatial and temporal resolutions, which are often not experimentally achievable. Various all-atom (AA) and coarse-grained (CG) protein force fields (FFs) have been developed, which differ in resolution, potential function, and parameterization strategy.^1–10^ The quality of the MD simulation results strongly depends on the accuracy of the FFs. Recent advances in computational power have allowed more extensive testing of AA and CG FFs with respect to experimental biophysical data, motivating researchers to refine these FFs further. Excessive aggregation of proteins was a common problem in various FFs.^11–14^ To address this issue, the NBFIX (NonBonded FIX) corrections to the AMBER and CHARMM FFs modified the interactions between amino acids using osmotic pressure data.^15–18^ MARTINI, the most widely used CG FF, also alleviated this problem by rebalancing nonbonded interactions in MARTINI 3.^19^ Another problem common to FFs focused on folded proteins is the limited accuracy of simulating intrinsically disordered proteins (IDPs), e.g., the dimensions of IDPs in aqueous solutions are too compact compared to experiments.^12,20,21^ Attempts have been made to optimize the interaction parameters, such as protein-water interactions, for both AA and CG FFs to achieve accurate simulations of IDPs.^12,21–23^ Compared to protein aggregations and IDPs, the accuracy of FFs in predicting the binding affinity of peripheral proteins to lipid membranes has not been extensively evaluated. However, a comparative study of the ability of AA FFs to reproduce the insertion energies of Wimley-White peptides at the membrane interface revealed excessive binding to some FFs.^24^

A CG protein model in SPICA FF^25–31^ has been recently developed, primarily targeting membrane proteins.^6,32^ In this model, the nonbonded parameters of side chains (SCs) were optimized to reproduce the hydration free energy of SC analogs, the potential mean force (PMF) between SC analogs in water, and the transfer free energy profile of SC analogs across dioleoyl phosphatidylcholine (DOPC) membranes. The nonbonded interactions of the backbones (BBs) were then determined by reproducing the penetration depth and tilt angle of the peripheral helices and the dimerization free energy of the transmembrane helices. The model developed using this parameterization protocol successfully simulated the stable structure of various membrane protein assemblies in DOPC bilayers and poliovirus capsids in aqueous solutions obtained from the Protein Data Bank (PDB) and AA-MD, respectively. However, this model has several problems similar to the other FFs mentioned above. For example, excessive membrane adsorption of peripheral proteins and too compact dimension of IDPs were observed.

In this study, we focused on improving the accuracy of the SPICA FF in simulating peripheral proteins and IDPs. First, we introduced different nonbonded Lennard-Jones (LJ) parameters for BB-water and BB-lipids, depending on the secondary structure of the proteins. In the previous model, the LJ parameters of the BBs were optimized against the reference data of α-helical peptides. However, peripheral proteins and IDPs contain large nonhelical regions, and the BB beads in these regions should be more polar than those in the helical regions of the CG model, which do not explicitly represent hydrogen atoms. Secondly, we re-parameterized interactions between amino acids, including alanine or charged SCs, which were overestimated in the previous protein model, leading to a non-negligible improvement. Finally, the protein-lipid interaction library was expanded via parameter optimization to reproduce the transfer free energy profiles of SCs through lipid membranes containing target lipids, including various glycerophospholipids, sphingolipids, and cholesterol, since the previous model only included phosphatidylcholine (PC). The new SPICA FF (SPICA FF ver2) successfully reproduced the radius of gyration (*R_g_*) of various IDPs, the binding sensitivity of several peripheral proteins, and the dimerization free energy of several transmembrane helices. The present model also showed improved performance with respect to the free energy of association of peptides in water. Furthermore, the specific binding of cholesterol to a G-protein coupled receptor (GPCR) was reproduced. These results suggest that CG MD simulations using SPICA FF ver2 provide highly accurate simulations for a wide range of protein systems.

## 2. METHODS

### 2.1 Coarse-grained model

The CG mapping of the 20 amino acids in the SPICA FF is shown in Figure S1. The BB atoms of one amino acid are represented as a single CG bead, and the SCs are composed of 0–4 CG beads, depending on their size and shape. Bond stretching, angle bending, and dihedral torsion interactions were described using simple potentials, as in the previous model.^6^ Coulombic electrostatic interactions were considered only between charged CG segments. LJ-type potential functions were employed to describe the nonbonded interactions.

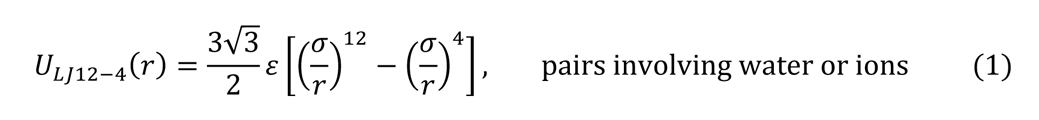

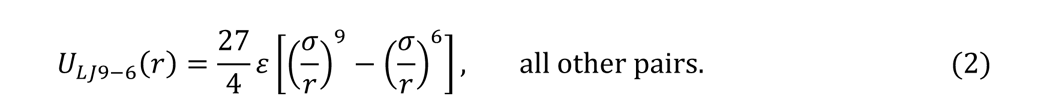

In the previous protein model in SPICA^6^, all pairs involving protein particles interacted via the LJ 9-6 potential (Eq. (2)). However, in the SPICA lipid model, the LJ12-4 potential (Eq. (1)) is used for pairs involving water or ions. Following the lipid model, we changed the potential function to LJ12-4 for the interactions between proteins and water/ions, as shown in Eq. (1). This allows for an intuitive comparison of the strength of the interactions of lipids and proteins with water in terms of hydrophobicity, which is advantageous when considering the strength balance of interactions. The LJ parameters of SC-water were re-optimized by reproducing the experimental hydration free energy of the SC analogs. To maintain the secondary structure of the proteins, an elastic network model (ENM) was applied with a force constant of 1.195 kcal/mol/Å^2^ to pairs of BB beads within 9 Å separated by three or more bonds, except in the case of IDP simulations.

### 2.2 Secondary structure-dependent backbone parameters

Secondary structure-dependent LJ parameters of the BB for water and lipids were introduced in the present model. Protein secondary structure assignments were made using the DSSP program^33^, which divides BB beads into three types: helix (α-helix, 3-helix, 5-helix), sheet (residue in isolated β-bridge, extended strand, hydrogen-bonded turn), and loop (bend, random coil). The secondary structure names are based on the definitions of the DSSP.

For the helix BB, the LJ parameters for BB-water were set to reproduce the hydration free energy of the BB beads calculated using the previous SPICA protein FF version^6^. In the previous model, the hydration free energy of the polypeptide helix obtained from the AA-MD was reference data for the BB-water interaction. Then, as in the previous protein model,^6^ the LJ parameters of helical BB-lipids were optimized by reproducing the penetration depths and tilt angles of various peripheral helical peptides at the DOPC membrane surface obtained from the Orientations of Proteins in Membranes (OPM) database.^34^ Finally, the nonbonded interactions of BB-BB and BB-SC were optimized to reproduce the dimerization free energy of the transmembrane helices.

The sheet BB bead was set to be slightly more hydrophilic than the helical BB to reproduce the *R_g_* of IDPs containing hydrogen-bonded turns. Then, we evaluated the penetration depths and tilt angles of β-hairpin peptides at the DOPC membrane surface, and compared with the OPM database^34^. For the loop BB, the LJ parameters for BB-water were optimized to reproduce the *R_g_* values of various IDPs in aqueous solutions. The interaction parameters between the loop BB and lipids were then optimized by reproducing the binding sensitivity of water-soluble and peripheral proteins. The LJ parameters for BB-BB and BB-SC in the sheet and loop regions were the same as those in the helix region. However, the good performance of the present model for the global dimension of IDPs and the association free energy of proteins in water indicates that the balance of interactions between BB-water and within proteins is reasonable (data shown in Section 3.1).

### 2.3 General setup for CG MD simulations

The initial configurations were built with an AA resolution using CHARMM-GUI^35^, and then mapped into the CG model using in-house tools (available at https://github.com/SPICA-group/spica-tools). The CG MD simulations were performed using the LAMMPS software package.^36^ Nonbonded LJ interactions were cut off at 15 Å, while electrostatic interactions were calculated using the particle-particle-particle-mesh (P3M) method.^37^ The energy minimization of the systems was conducted using the conjugate gradient algorithm, and the production MD runs were carried out with a time step of 10 fs in the NPT ensemble. The Nosé–Hoover thermostat^38,39^ and Parrinello-Rahman barostat^40,41^ were used to control the temperature and pressure, respectively. The pressure was maintained at 1 atm, and the target temperature was changed slightly for each system.

### 2.4 Calculating the dimerization free energy of peptides

The dimerization free energy, Δ*G*_)*+_, of transmembrane peptides in lipid bilayers (2-dimensional) was calculated using the following equations^19^:

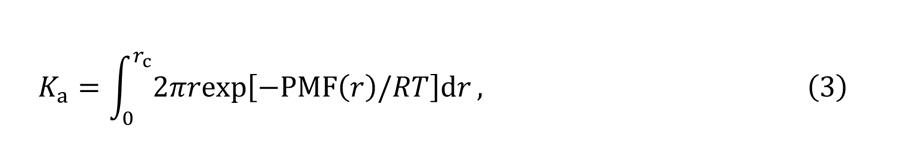

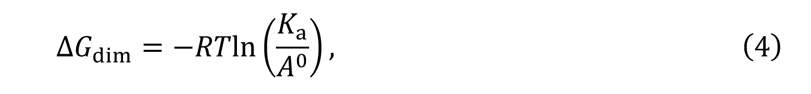

where *r* and PMF(*r*) are the distances between the center of mass (COM) of the monomer peptides on the membrane and the free energy profile along *r*, respectively, and *r*_/_ is the cut-off distance in the integration, which was set to 35 Å. The standard area Å. was set to 100 Å^2^, which was used in the FRET experiments.^42^ The PMF shifted to zero in the plateau region before integration.

The Δ*G*_)*+_of peptides in an aqueous solution (3-dimensional) was calculated using the following equations^43^:

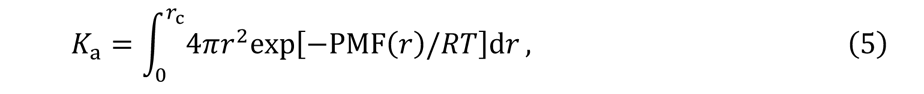

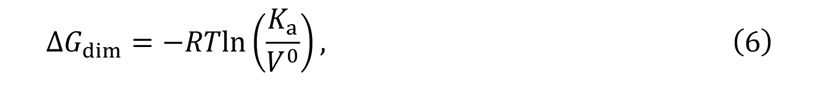

where *r* and PMF(*r*) are the distances between the COM of the monomer peptides in water and the free energy profile along *r*, respectively. The cut-off distance *r*_/_ and standard volume *V*^0^ were set to 45 Å and 1661 Å^3^, respectively.

The dimerization free energies of the transmembrane helices were calculated from the PMF using Eq. (3) and (4). For the PMF calculations, two monomers of GpA, WALP23, and SerZip, and the transmembrane segments of EphA1 and ErbB1 were inserted into the DPPC, DOPC, POPC, DMPC, and DLPC bilayers, respectively. Each bilayer membrane is composed of 256 lipid molecules and abundant water to obtain a fully hydrated membrane. Two WALP23 monomers were placed in an antiparallel configuration, whereas the other peptides were placed in parallel configurations. The systems of EphA1 and ErbB1 were ionized at NaCl concentrations of 0.15 M and 0.50 M, respectively. The systems were then neutralized by adding Na^+^ or Cl^-^ ions as needed.

The PMFs were calculated using the adaptive biasing force (ABF) method.^44^ The reaction coordinate *r* was split into six windows spanning from 3 to 35 Å. To match the experimental conditions, the temperatures were set to 323, 308, 310, 303, and 303 K for GpA, WALP23, SerZip, EphA1, and ErbB1, respectively. ABF sampling was performed by running a 4-μs MD simulation for each window of each system. The averages and standard deviations of dimerization free energies were estimated from the block data obtained from each 1 μs of sampling.

The dimerization free energies of the A3-B3 and insulin peptides were calculated from the PMF using Eqs. (5) and (6), respectively. A3-B3 peptides are artificially synthesized coiled-coil peptides^45^ and the structures of the A3 and B3 peptides were modeled as ideal α-helices (A3: GEIAALEKENAALEWEIAALEQGG, B3: GKIAALKYKNAALKKKIAALKQGG). Two insulin monomers (PDB ID: 4INS) were put in water with a box length of 105 Å, while the A3-B3 peptides were placed in a box with a length of 120 Å. The system containing A3-B3 peptides was ionized at a concentration of 0.14 M NaCl. The systems were then neutralized by adding Na^+^ or Cl^-^ ions. The PMF was estimated using the ABF method. The reaction coordinate *r* was the distance between the COMs of the two monomers, divided into six windows ranging from 15 to 45 Å for the insulin monomers and 3 to 45 Å for the A3-B3 peptides. The temperatures were set at 300 and 293 K for insulin and A3-B3 peptides, respectively, according to the experimental conditions. MD was performed for 3 and 5 μs for each window of the insulin and A3-B3 peptides, respectively.

### 2.5 IDP simulations

To optimize the LJ interaction parameters between the BB segment of the protein loop and sheet region and water, we evaluated the *R_g_* values of 10 IDPs over a wide range of experimental *R_g_* values. We selected 9 IDPs from the previous study^21^: Histatin-5 (Hst5), a two repeat of Histatin-5 (Hst52), the activation domain of ACTR (ACTR), the T-domain of colicin N (ColNT), the low-complexity domain of hnRNPA1 (A1), α-synuclein (aSyn), the plug domain from a TonB-dependent receptor (FhuA), and two deletion mutants of Tau (K19 and K25). The Mengo virus leader protein (Leader)^46^ was added to this set. The IDPs were solvated and ionized at the salt concentrations used in the experiments (Table S1). The ENM was applied only within the structured regions. MD simulations were run for 1 μs at the temperatures of the experimental conditions (Table S1) for two replicas of each system; different initial velocities were given for the two replicas. The *R_g_* values were then calculated and standard errors were estimated using block analysis. The first 300 ns of the MD trajectories of the two replicas were not used in the *R_g_* calculation. Information on the simulation systems and experimental *R_g_* values for each IDP is presented in Table S1.

### 2.6 Calculations of the free energy profiles of SC analogs across lipid bilayers

To optimize the interactions between SCs and various lipid species, the free energy profile of SC analogs across lipid membranes were evaluated. A bilayer system, including one target SC analog, was built and neutralized with Na^+^ and Cl^-^ when needed. Detailed system information is presented in Table S2. The free energy profile was calculated using the ABF method.^44^ The reaction coordinate was the projected distance to the bilayer normal (*z*-direction) between the COMs of SC analogs and the lipid bilayer, divided into six windows ranging from 0 to 30 Å at 5-Å intervals. A 100-ns CG MD run, which proved to be sufficient for convergence in our previous study^6^, was performed for each window of each system. We evaluated the same from the AA-MD using the CHARMM36 FF to obtain reference data; in this case, a 300–600-ns AA-MD run was required for each window to observe the convergence of the free energy profiles.

### 2.7 Simulations of protein adsorption to lipid membranes

To optimize the LJ parameters between BBs and lipids, we evaluated the binding sensitivity of proteins to the lipid membranes from CG MD simulations. To do so, the protein was placed in water region, away from the membrane such that the minimum distance between the protein and lipid molecules was at least 15 Å. Four replicates of the simulations with different initial velocities were performed for each system. A 2-μs MD simulation was performed at 310 K for the four replicates of each system, except for the aSyn system. For the aSyn system, a 1-μs MD simulation was run for four replicates at a temperature of 283 K.

We used a previously proposed method to quantify membrane binding by proteins.^47^ Time traces of the minimum distances between the proteins and lipid molecules were calculated from the CG MD simulations, and normalized probability distributions of minimum distances were constructed using kernel density estimation. The bound state was defined as the state when the minimum distance was smaller than 7 Å, and the bound fraction was calculated as the integral of the distribution curve up to a distance of 7 Å. Averages and standard deviations of the bound fractions were computed from four replicates.

To detect the membrane-binding residues of the protein from the CG MD simulations, the normalized contact frequencies of the residues were computed using the following protocol: a residue was regarded as binding to the membrane if the distance between any bead of the residue and any lipid bead was smaller than 5 Å. For each residue, the number of binding events during the MD trajectory was counted and summed over all replicates to yield the normalized contact frequency.

## 3. RESULTS AND DISCUSSION

Because the LJ function for protein-water interactions was changed, the LJ parameters between SCs and water were adjusted to reproduce the experimental values of the hydration free energy of the SC analogs, as was done in the previous SPICA FF. Similarly, as shown in Table S3, good optimization was achieved to obtain quantitative agreement with the experiments.

Note that the optimization of the amino acid nonbonded parameters shown in the following subsections has been performed several times, and all results from SPICA ver2 were obtained with the same final parameter set.

### 3.1 Protein-protein interactions and global dimension of IDPs

#### 3.1.1 Protein-protein interactions

The previous SPICA FF tended to overestimate some interactions involving charged SCs or alanine. Here, we optimized these interaction parameters by more accurately reproducing the radial distribution function (RDF) of pairs involving charged SCs in water, as calculated from AA-MD simulations, and the dimerization free energy between five transmembrane helices (GpA, WALP23, SerZip, and the transmembrane segments of EphA1 and ErbB1). The free energies of association of the two insulin monomers and A3-B3^45^ in water were then evaluated using CG MD simulations and compared with the experimental data. Insulin is a relatively small peptide (51 amino acids) and includes helix, sheet, and loop regions. The A3-B3 peptide contains many charged and alanine residues. Hence, these peptides are suitable for validating interaction parameters in the improved SPICA FF.

The RDFs between CG segments containing charged SCs were calculated from the MD trajectories of connected SC dimers in aqueous solutions, as in our previous study.^6^ The RDFs were calculated using the CHARMM FF, the previous SPICA FF, and the present SPICA FF (Figure S2). The previous SPICA FF (ver1) occasionally gave a higher first peak of the RDF than the CHARMM FF, indicating that these pairs were too adhesive in aqueous solutions. This was particularly pronounced in the pairs involving Arg SCs. Overall, the excessive height of the first peak of the RDF observed in the previous SPICA FF was suppressed in the present SPICA FF, indicating that the present SPICA FF provides moderate association properties comparable to those of the AA model between SC dimers in aqueous solutions.

The Δ*G*_)*+_ of the five transmembrane helices in lipid membranes and the two peptides in water, as obtained from CG MD simulations, are plotted in Figure 1, along with the experimental values. For the transmembrane helices, the Δ*G*_)*+_ calculated in the previous SPICA FF showed good agreement with those obtained from experiments, except for WALP23, an alanine-rich peptide. Excessive binding of the WALP23 peptides observed in the previous version was successfully improved, primarily by modifying the interactions between the alanine segments in the SPICA FF (Figure 1a). For the other transmembrane peptides, the Δ*G*_)*+_values calculated via MD simulations using the current SPICA FF (ver2) were in quantitative agreement with the experimental estimates. Furthermore, the association free energies in water between the A3-B3 peptides and between two insulin peptides calculated using the current SPICA FF were in reasonable agreement with the experiments (Figure 1b), in contrast to the previous SPICA FF, which showed excessive association for these two peptides. The improvement in the dimerization free energy was achieved by improving the pair-interaction parameters involving charged SCs or alanine segments and, in part, by re-optimizing the interaction parameters of the BB segments depending on the protein secondary structure.

**Figure 1.**
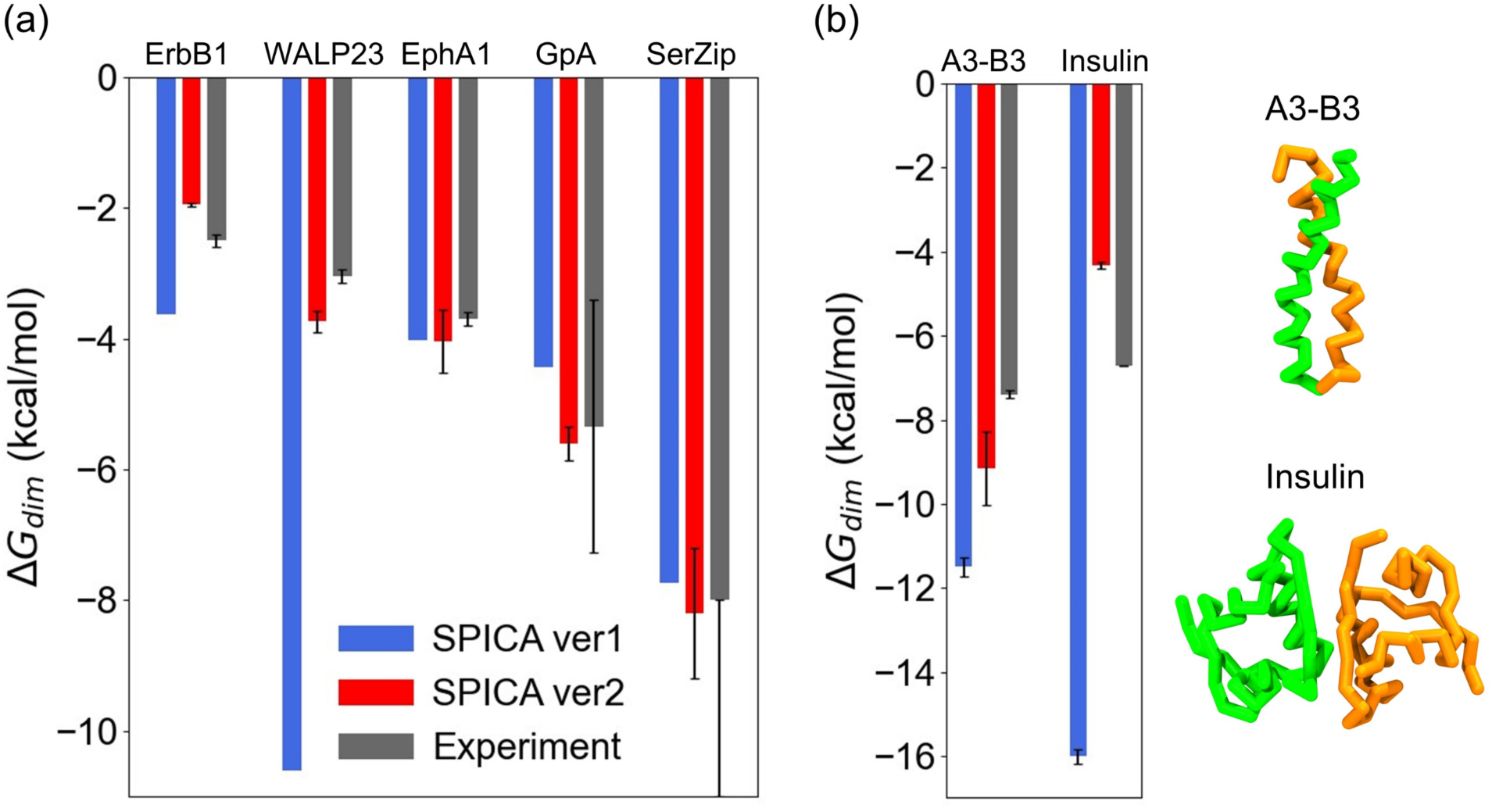
Dimerization free energies of (a) five transmembrane helices in lipid membranes and (b) two peptides in an aqueous solution. Experimental data were obtained from the literature.^45,48−^ _53_ The experimental data for SerZip was less than −8 kcal/mol (more negative).^52^

#### 3.1.2 Global dimension of IDPs

Figure 2a plots the *R_g_* values of 10 IDPs obtained from CG MD against the experimental *R_g_*. The MD calculations using the previous SPICA FF (ver1) were performed only for the Hst5 and aSyn systems, and their *R_g_* values were much smaller than those obtained from the experiments; in particular, the average *R_g_* of aSyn was approximately 40 % of the experimental value. In the MD calculations using the present SPICA FF (ver2), the *R_g_* values were greatly improved and agreed well with the experimental values, except for A1 and K19. For A1 and K19, the *R_g_* values still showed relatively large deviations of approximately +30 % and −20 % compared to the experimental values, respectively. The good performances for the dimension of IDPs in water and protein associations in aqueous solution and lipid membranes indicate that the interactions of proteins with water and lipids are well balanced in the present SPICA FF ver2.

**Figure 2.**
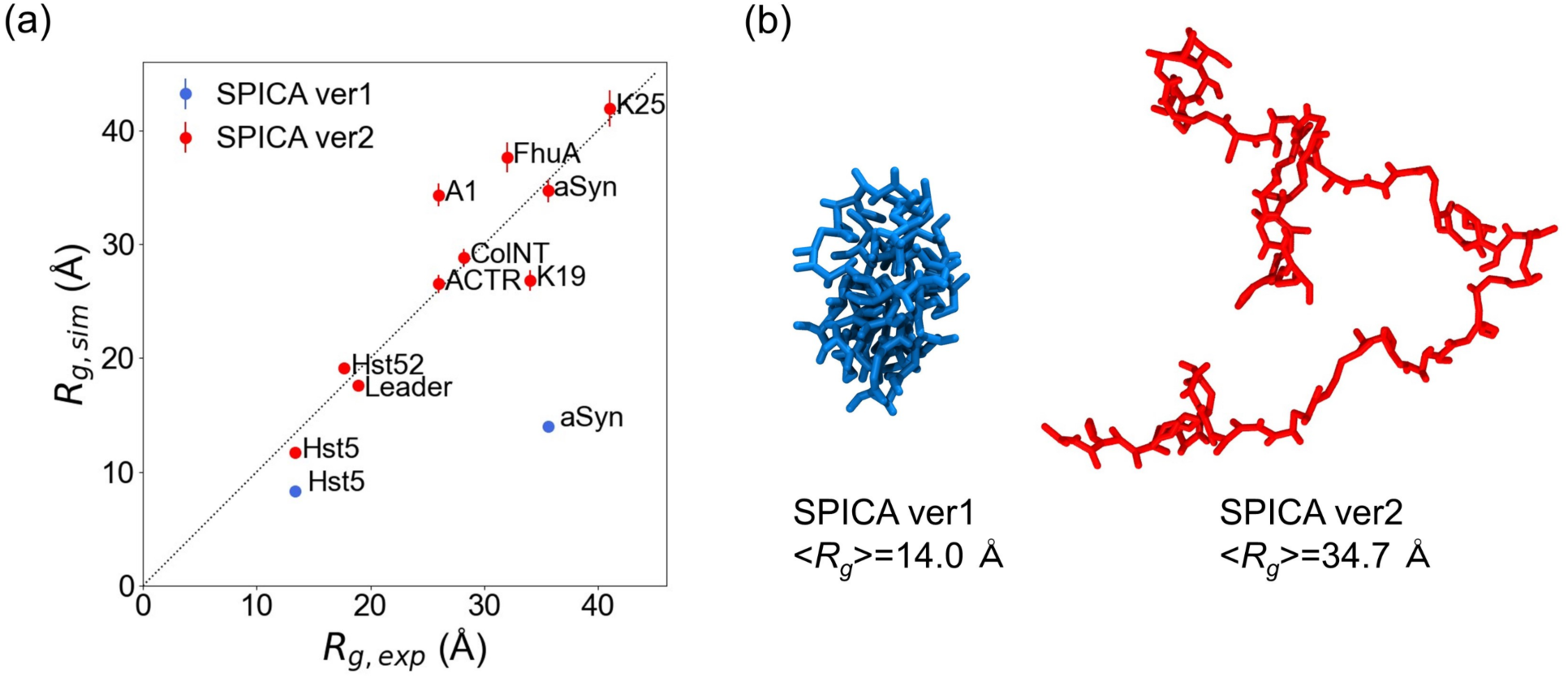
(a) *R_g_* values calculated from CG MD are plotted against the experimental *R_g_* values. The dotted line indicates perfect agreement between CG MD simulations and the experiments. (b) Typical snapshots of aSyn at the average *R_g_* obtained from the CG MD using SPICA ver1 (in blue) and ver2 (in red). Solvent molecules are omitted for clarity.

### 3.2 Expanding library of protein-lipid interactions

#### 3.2.1 Free energy profiles of SC analogs across various lipid membranes

Since the LJ interaction function between protein segments and water has been changed to LJ12-4, the LJ parameter (*ε*) of the SC analogs with respect to lipid segments needs to be re-optimized.

The free energy profiles of the SC analogs to pass through lipid membranes were used for this purpose. In the previous SPICA study, we optimized the SC parameters only with the DOPC membrane. In the present study, we evaluated the free energy profiles of SC analogs across lipid bilayers to optimize the interactions between SCs and various other lipid species. The target lipids for optimization, in addition to DOPC, included cholesterol, sphingomyelin (SM), and four 1-palmitoyl-2-oleoyl-glycerophspholipids, which have different head species: e.g., ethanolamine (PE), glycerol (PG), inositol (PI), and serine (PS).

The free energy profiles of SCs across the 1-palmitoyl-2-oleoyl-phosphatidyl serine (POPS) bilayer, calculated using CG MD with SPICA FF ver2 and AA-MD, are shown in Figure 3 (see Figures S3–S8 for the other lipid bilayer systems). Although we previously used the reference free energy profiles obtained from AA-MD with the OPLS-AA FF^3^, we computed the reference free energy profiles using CHARMM36 FF^1^ in this study to maintain consistency with the other reference data used in the SPICA FF. The free energy profiles of all SCs across the POPS bilayer obtained from SPICA FF ver2 showed excellent agreement with those obtained from AA-MD with the CHARMM FF, as shown in Figure 3. The same was true for all the pure phospholipid bilayer systems (Figs. S3–S6); however, a degraded match was detected for membranes containing cholesterol and SM (Figs. S7 and S8). In particular, the free energy profiles of charged SCs crossing these lipid bilayers with higher-order hydrophobic cores were less accurately described using the present SPICA FF. This limitation most likely originates from the entropy in the membrane core, which may not be easily corrected in the CG description.

**Figure 3.**
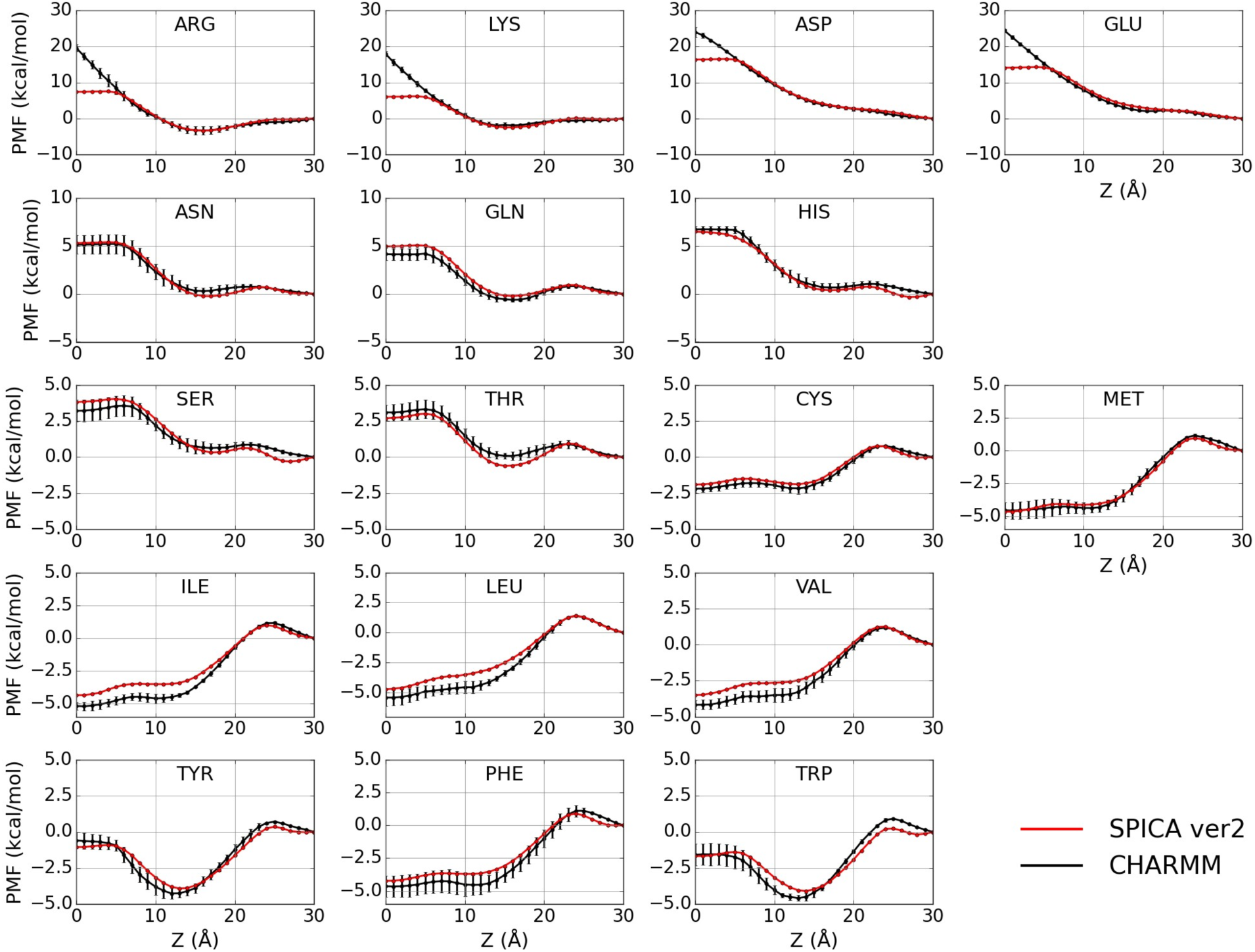
The transfer free energy profile of SC analogs across the POPS bilayer as calculated from the CG and AA-MD simulations. The origin corresponds to the center of bilayers and 25–30 Å corresponds to the bulk water region.

#### 3.2.2 Cholesterol-binding sites in G-protein coupled receptor

Cholesterol is known to play a crucial role in regulating the functions of various GPCRs.^54^ The specific binding of cholesterol affects the structural stability of GPCRs.^55,56^ Here, we investigated whether the modified lipid-protein interactions in SPICA FF ver2 could reproduce cholesterol binding to the experimentally observed binding site. We performed a CG MD simulation with the β_2_-adrenergic receptor (β_2_AR, PDB ID: 3D4S)^57^ inserted into a bilayer consisting of 170 POPC and 84 cholesterol molecules under the same conditions as in a previous AA-MD study.^58^ A MD simulation was performed with a temperature of 310 K for 3 μs.

The binding of cholesterol to the binding site in the crystal structure was observed (Figure 4a). The interaction between the hydroxyl group of cholesterol and the basic residues (R/K) of the receptors is a common feature of cholesterol-binding sites.^59^ This interaction was successfully reproduced using SPICA FF ver2 (Figure 4b). We also analyzed the spatial distribution of cholesterol molecules around β_2_AR (data not shown). The extremely high probability of cholesterol molecules being present at the binding site indicated the specific binding of cholesterol. These results demonstrate the ability of SPICA FF ver2 to capture specific interactions between cholesterol and receptor proteins.

**Figure 4.**
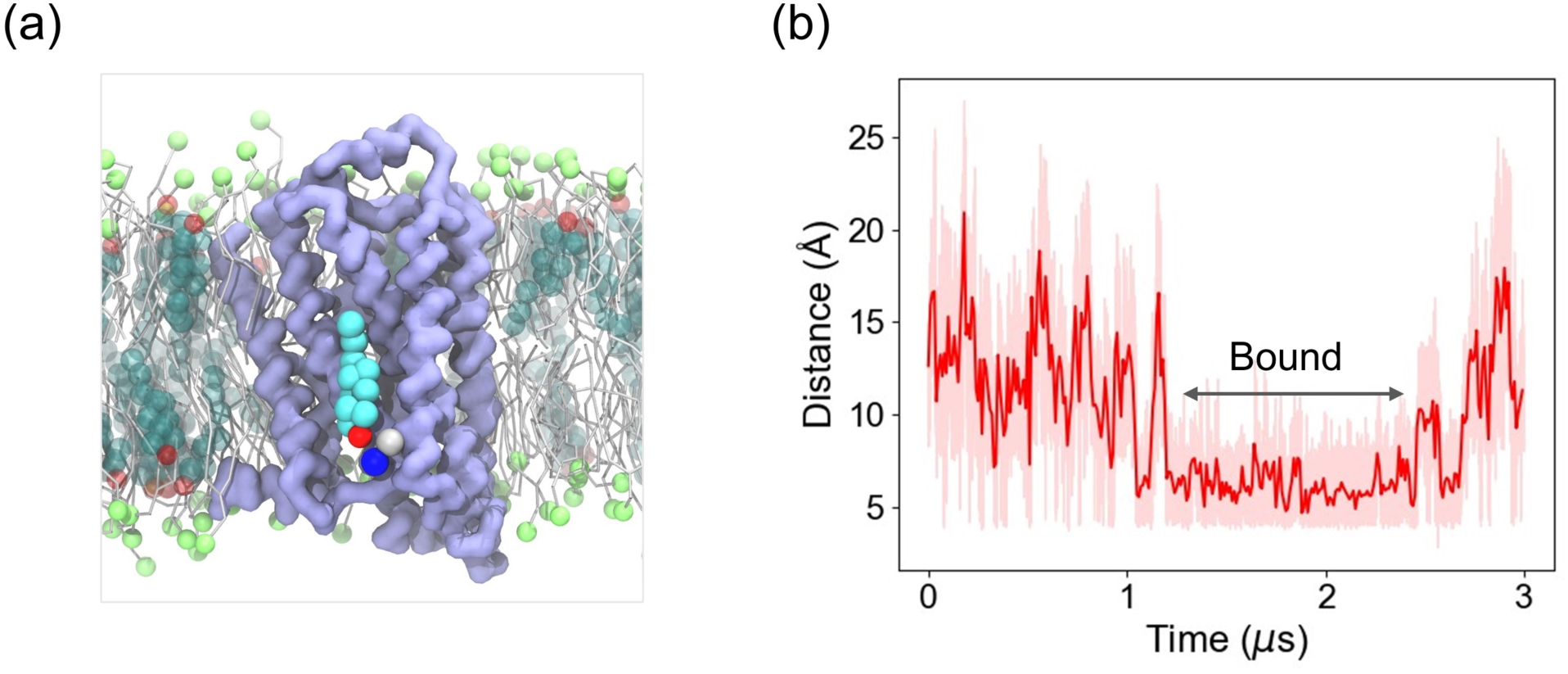
Specific binding of cholesterol to the β2AR. (a) Snapshot of a cholesterol molecule in the binding site. β2AR is shown in light blue. The AR1 and AR2 beads of the Arg SC in the binding site are represented as VDW spheres and shown in silver and blue, respectively. The hydroxyl groups and other parts of the cholesterol molecule are represented as VDW spheres and shown in red and cyan, respectively. A cholesterol molecule in the binding site is represented as opaque, while the other cholesterol molecules are shown as transparent. The head groups and other parts of the POPC molecule are represented as lime VDW spheres and silver licorice, respectively. Solvent molecules are omitted for clarity. (b) Time trace of the minimum distances between the hydroxyl groups of all cholesterol molecules (red in Figure 4a) and the AR2 particle in the binding site (blue in Figure 4a). The shaded line corresponds to the raw data; the solid line is the block average of the data.

### 3.3 Protein-membrane interactions

The previous SPICA FF tended to over-stabilize the adsorbed state of proteins on the surface of lipid membranes. For example, proteins known to remain soluble in the cytosol strongly bound to the DOPC membranes. In the present study, we modified the LJ parameters of BB-lipids by qualitatively reproducing the binding sensitivity of proteins to lipid membranes. To this end, we checked the binding of four proteins: lysozyme, phospholipase A_2_ (PLA_2_), the pleckstrin homology (PH) domain of 3-phosphoinositide-dependent kinase-1 (PDK1), and α-synuclein (aSyn). Lysozyme is known to remain soluble in the absence of negatively charged lipids^60^, PLA_2_ has an affinity for the DOPC membrane^61^, the PH domain of PDK1 is sensitive to PS lipids^62^, and aSyn binds to the membrane, mimicking the composition of a synaptic vesicle.^63^ aSyn fully adopts the random coil structure in water, while the membrane-interacting region (residues 1–98) folds into a helical conformation at the membrane interface. In the present study, we used a micelle-bound aSyn structure (PDB ID: 1XQ8)^64^ and applied ENM only to the helical regions. The composition of the lipid membrane in the CG MD simulations and the binding sensitivities of these proteins are summarized in Table 1.

**Table 1.** System information for the protein-membrane interactions and the bound fractions calculated from CG MD simulations.

| Protein | PDB ID | Lipid composition | Binding sensitivity from experiments | Bound fraction from CG MD (%) |
| --- | --- | --- | --- | --- |
| Lysozyme | 1AKI | 400 DOPC | Water soluble | $4.4 \pm 1.1$ |
| PLA<sub>2</sub> | 1POA | 400 DOPC | Binding to DOPC membrane | $67.2 \pm 24.0$ |
| PDK1 PH domain (residues 407–549) | 1W1D | 400 DOPC | Sensitive to PS lipid | $7.7 \pm 2.6$ |
| | | 320 DOPC, 80 DOPS | | $56.4 \pm 23.7$ |
| aSyn | 1XQ8 | 940 DOPE, 564 DOPS, 376 DOPC (50:30:20) | Binding to the membrane | $88.0 \pm 19.3$ |

The bound fractions calculated from the CG MD simulations for each system are listed in Table 1. SPICA FF ver2 successfully reproduced the non-binding of lysozyme to DOPC membranes (average bound fraction: 4.4 %) and the binding of PLA_2_ to DOPC membranes (average bound fraction: 67.2 %). The membrane-interacting residues of PLA_2_ were analyzed by calculating the normalized contact frequency of each residue with the membrane. The experimentally observed binding interface at which the aromatic residues of PLA_2_ (Y3, W18, W19, W61, F64, and Y110) interact with lipids^61,65^ was reproduced using SPICA FF ver2 (Figure 5a and 5d). The PS sensitivity of the PDK1 PH domain was also reproduced, as shown in Table 1 (average bound fractions of 7.7 % for DOPC and 56.4 % for DOPC/DOPS). Membrane-interacting residue analysis of the PDK1 PH domain captured binding by the experimentally observed binding residues R466 and K467 (Figure 5b and 5e)^62^, which strongly interact with the negatively charged PS lipids. The binding of aSyn to the membrane, mimicking the composition of synaptic vesicles, was also reproduced using SPICA FF ver2 (average bound fraction: 88.0 %). Furthermore, the experimentally proposed binding mode was also reproduced, with the helical region (residues 1– 98) binding to the membrane and the loop region (residues 99–140) interacting weakly with the membrane^63^ (Figure 5c and 5f). These results indicate that the present SPICA FF ver2 can be used to evaluate the membrane-water partitioning (binding ratio) of peripheral proteins and determine the membrane-interacting residues of proteins.

**Figure 5.**
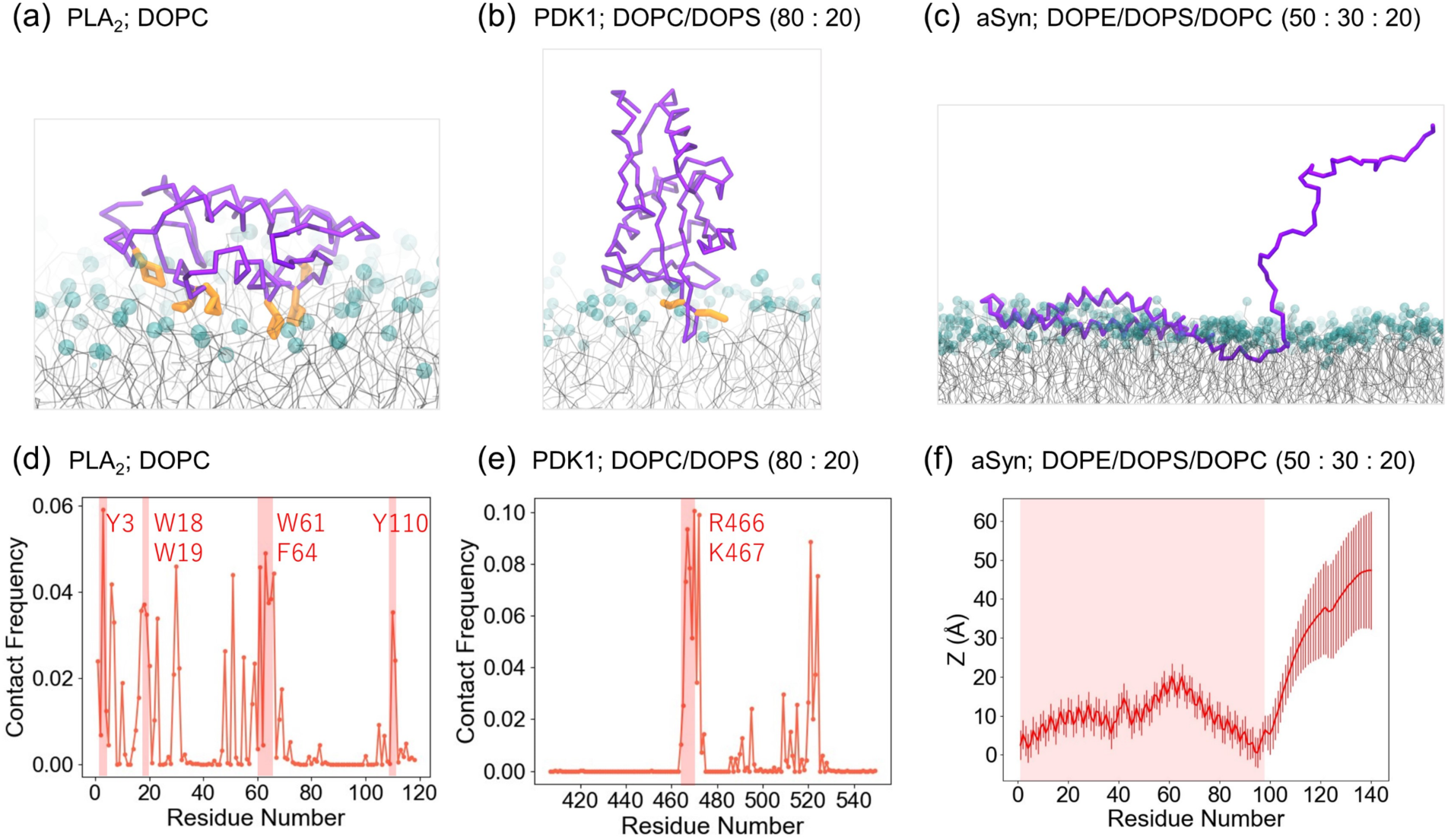
Protein binding modes are shown as snapshots of (a) the PLA2-DOPC system, (b) the PDK1-DOPC/DOPS system, and (c) the aSyn-DOPE/DOPS/DOPC system. The backbone (BB) beads of the proteins are shown in purple. The head groups and the rest of the lipids are represented as cyan VDW spheres and gray bonds, respectively. The experimentally determined residues involved in membrane binding are shown in orange. (d, e) Normalized contact frequency of the protein residues with the membranes. Shaded regions represent experimentally observed binding residues. (f) Average *z*-distance of the BB beads for each residue after the adsorption of aSyn to the lipid membrane from the average lipid phosphate position in the upper leaflet of the bilayer. A shaded region represents the experimentally observed binding region.

Here, the LJ parameters of the BB-lipid interactions were modified to obtain the proper adsorption behavior of some proteins on lipid membranes. In our previous model, the LJ parameters of BB-lipids were optimized using the OPM database^34^ against the penetration depth and tilt angle of the peripheral helices on the DOPC membrane surface. A series of test MD simulations were performed to verify the performance of the SPICA FF against the OPM database. A comparison with the reference data (OPM database) still showed good agreement for the *α*-helices (Figure S9) and was in reasonable agreement with the OPM database for the β-hairpin peptides (Figure S10).^6^ Hence, we succeeded in improving the adsorption behavior of proteins in SPICA FF ver2 without compromising accuracy of the agreement with the OPM database.

## 4. CONCLUSIONS

In this study, we propose an improved protein model in SPICA FF by introducing a secondary structure-dependent LJ parameter for protein BB segments and rebalancing most interactions involving the proteins. The LJ parameters of BB beads were optimized to reproduce various target properties of peptides and proteins, such as the penetration depth and tilt angle of peripheral peptides, the dimerization free energy of transmembrane helices, the *R_g_* of IDPs, and the binding sensitivity of peripheral proteins. The new model, SPICA FF ver2, also showed reasonable performance for peptide association free energies in water. These results shows that the nonbonded interactions involving proteins are well balanced in SPICA FF ver2. In addition, owing to the optimized interactions between proteins and various lipids, SPICA FF ver2 succeeded in reproducing the selective binding of several peripheral proteins by their correct binding interfaces and the specific interactions of cholesterol molecules with the receptor protein. Overall, the SPICA FF ver2 is applicable with high accuracy for simulating extensive protein systems. Specifically, our new strategy of using secondary structure-dependent LJ parameters for protein BB segments, overcomes known limitations of existing “pragmatic” FF^66^ without the need of specific ad-hoc rescaling or parameterization strategies^21,67,68^. Our work suggests that there is still ample room for improvement towards the accuracy of chemically-transferable pragmatic CG force fields using chemistry-driven intuition.

## ACKNOWLEDGMENTS

This study was supported by the Japan Society for the Promotion of Science (JSPS) KAKENHI (Grant number JP 21H01880 to WS) and by the Swiss National Supercomputing Centre (CSCS) (Project ID s1221 to SV). This study used the computational facilities of the Research Center for Computational Science, Okazaki, Japan (Project:22-IMS-C108, 23-IMS-C095) and the Institute for Solid State Physics, University of Tokyo, Japan. SV and AK acknowledge support by the Swiss National Science Foundation through the National Center of Competence in Research Bio-Inspired Materials.

## DATA AVAILABILITY

All parameters and simulation tools used to prepare the initial setups for the MD simulations are freely available on the webpage (https://www.spica-ff.org/) and github (https://github.com/SPICA-group/spica-tools), respectively.

## ABBREVIATIONS

GpA, glycophorin A; SerZip, serine zipper; DPPC, 1,2-dipalmitoyl-sn-glycero-3-phosphocholine; POPC, 1-palmitoyl-2-oleoyl-glycero-3-phosphocholine; DMPC, 1,2-dimyristoyl-sn-glycero-3-phosphocholine; DLPC, 1,2-dilauroyl-sn-glycero-3-phosphocholine.

