## Supporting Information for "Improved Protein Model in SPICA Force Field"

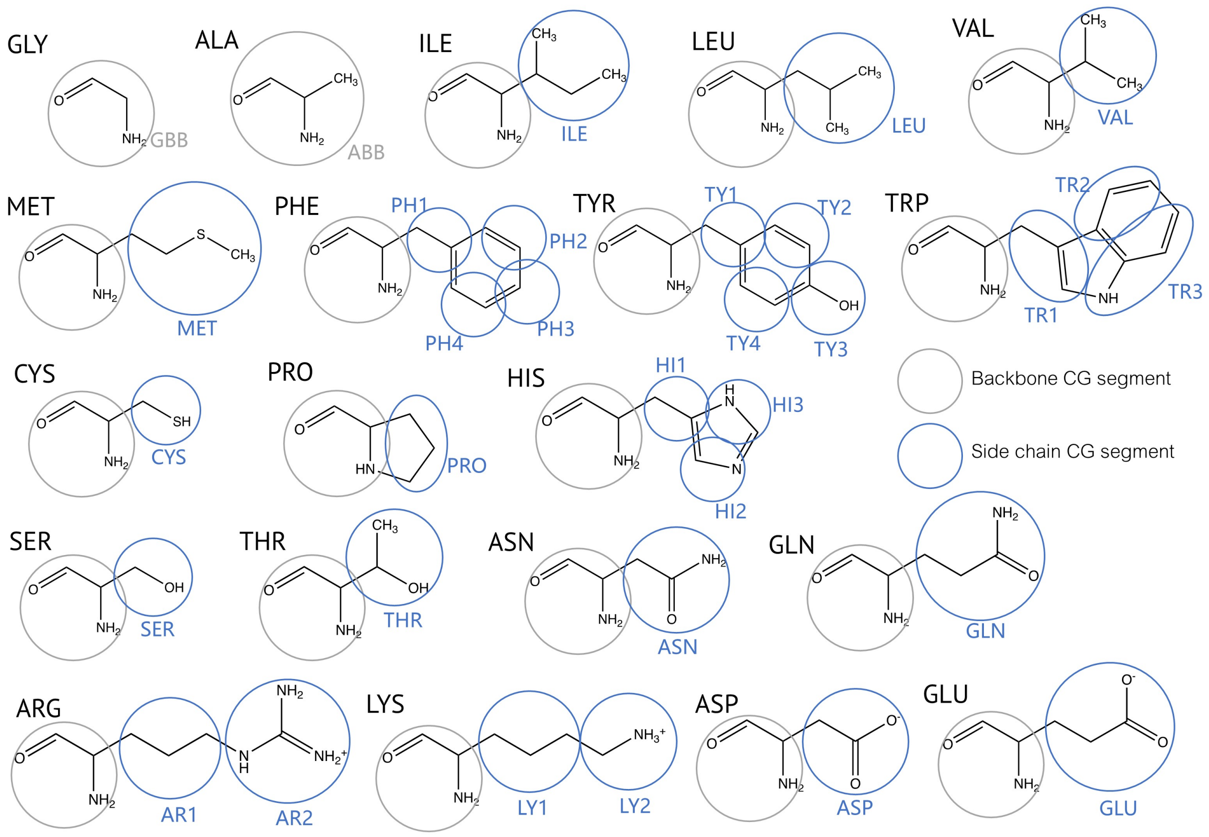


**Figure S1.** CG mapping of 20 amino acids in the SPICA FF.

**Table S1.** System details for IDP simulations: Number of amino acids residues ($N_R$), simulation temperature ($T$), box size ($d$), and salt concentration in the simulation ($C_s$)

| protein | $N_R$ | Exprimental $R_g$ [Å] | $T$ [K] | $d$ [Å] | $C_s$ [M] |
|---|---|---|---|---|---|
| Hst5 | 24 | 13.4 ± 0.5 | 293 | 80 | 0.15 |
| $(Hst5)_2$ | 48 | 17.7 ± 0.5 | 298 | 180 | 0.15 |
| Leader | 67 | 18.9 ± 2.0 | 277 | 130 | 0.15 |
| ACTR | 71 | 26.0 ± 3.0 | 278 | 190 | 0.2 |
| ColNT | 98 | 28.2 ± 0.3 | 277 | 210 | 0.4 |
| K19 | 99 | 34.0 ± 3.0 | 288 | 215 | 0.15 |
| A1 | 137 | 26.0 ± 1.0 | 296 | 215 | 0.05 |
| aSyn | 140 | 35.6 ± 0.4 | 293 | 240 | 0.2 |
| FhuA | 144 | 32.0 ± 2.0 | 298 | 220 | 0.15 |
| K25 | 185 | 41.0 ± 3.0 | 288 | 255 | 0.15 |

**Table S2**. System information for the calculations of free energy profile of SC analogues across the lipid bilayers.

| Target lipid | Lipid composition | Temperature (K) |
|---|---|---|
| DOPC | 72 DOPC | 310 |
| POPE | 72 POPE | 310 |
| POPG | 72 POPG | 310 |
| POPS | 72 POPS | 310 |
| POPI | 72 POPI | 310 |
| Cholesterol (CHOL) | 50 DPPC, 22 CHOL | 310 |
| Sphingomyelin(SM) | 128 stearoyl SM (SSM) | 323 |

**Table S3**. The hydration free energy of SC analogues from CG MD and experiments.

| Residue | SC analogue | $\Delta G_{hyd}$ (kcal/mol) | | Residue | SC analogue | $\Delta G_{hyd}$ (kcal/mol) | |
|---|---|---|---|---|---|---|---|
| | | SPICA | Exp. | | | SPICA | Exp. |
| LEU | Isobutane | 2.46 | 2.28 | THR | Ethanol | −4.70 | −4.88 |
| ILE | Butane | 2.26 | 2.15 | SER | Methanol | −5.00 | −5.06 |
| VAL | Propane | 2.12 | 1.99 | TRP | 3-methylindole | −5.84 | −5.88 |
| PHE | Toluene | −0.74 | −0.76 | TYR | Cresol | −6.08 | −6.11 |
| CYS | Methanethiol | −1.24 | −1.24 | GLN | Propionamide | −9.34 | −9.38 |
| MET | Methyl-ethyl-sulfide | −1.37 | −1.48 | ASN | Acetamide | −9.61 | −9.68 |

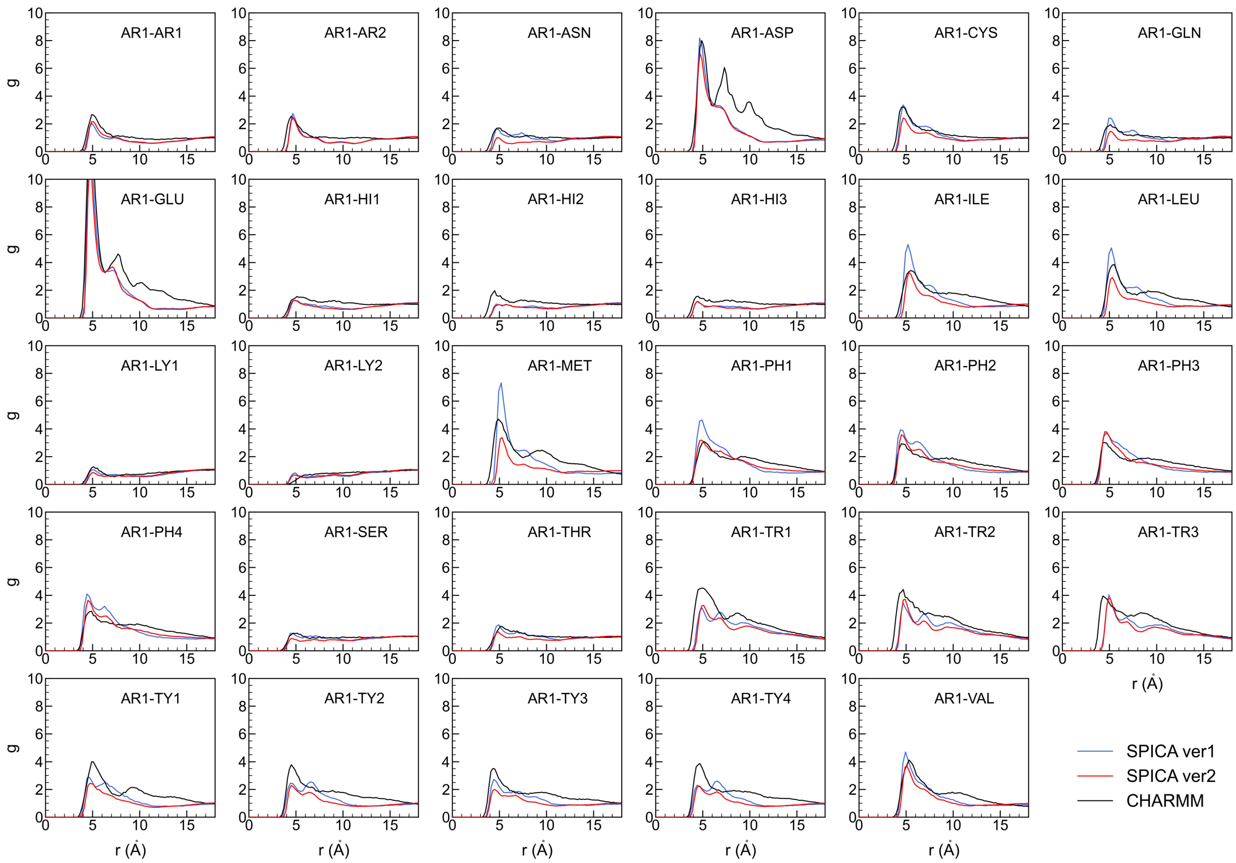
AR1-AR1
AR1-AR2
AR1-ASN
AR1-ASP
AR1-CYS
AR1-GLN
AR1-GLU
AR1-HI1
AR1-HI2
AR1-HI3
AR1-ILE
AR1-LEU
AR1-LY1
AR1-LY2
AR1-MET
AR1-PH1
AR1-PH2
AR1-PH3
AR1-PH4
AR1-SER
AR1-THR
AR1-TR1
AR1-TR2
AR1-TR3
AR1-TY1
AR1-TY2
AR1-TY3
AR1-TY4
AR1-VAL
g
r (Å)
SPICA ver1
SPICA ver2
CHARMM

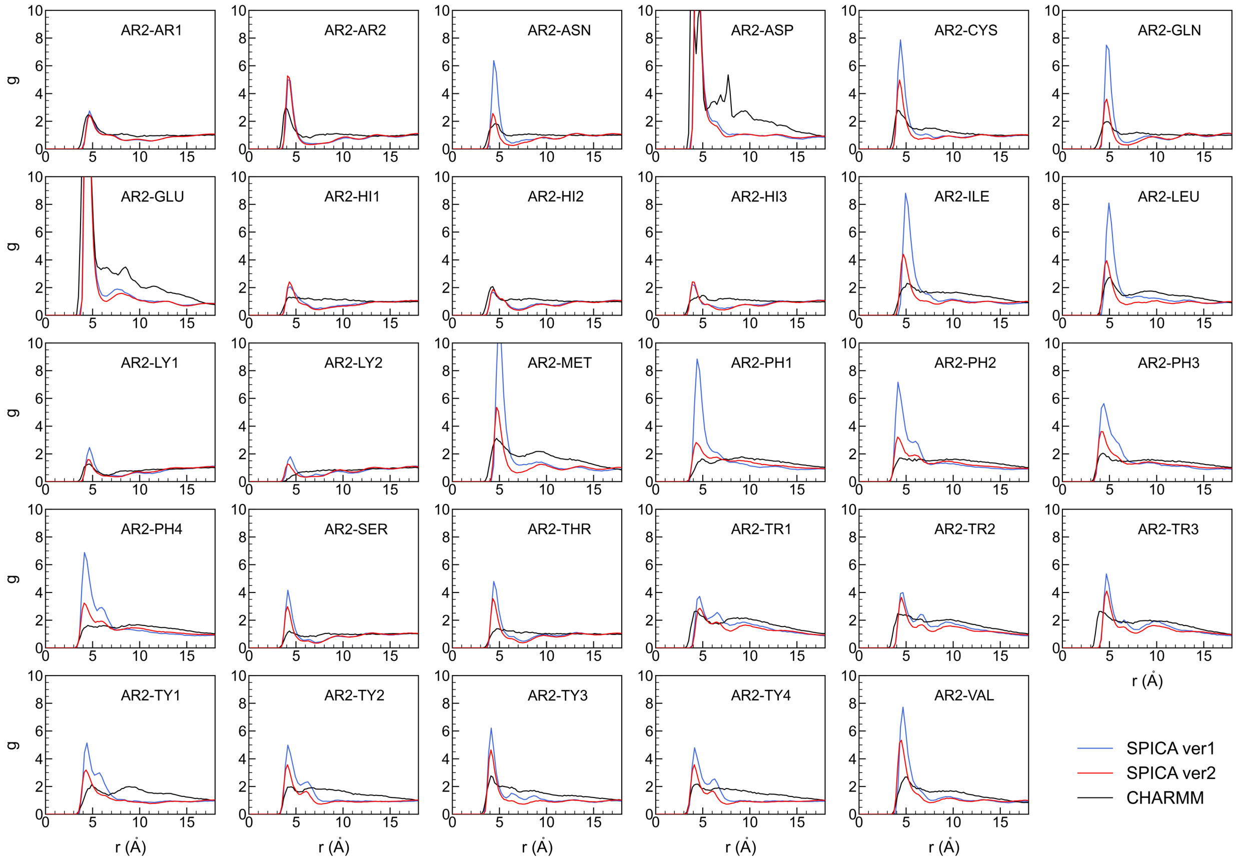
AR2-AR1
AR2-AR2
AR2-ASN
AR2-ASP
AR2-CYS
AR2-GLN
AR2-GLU
AR2-HI1
AR2-HI2
AR2-HI3
AR2-ILE
AR2-LEU
AR2-LY1
AR2-LY2
AR2-MET
AR2-PH1
AR2-PH2
AR2-PH3
AR2-PH4
AR2-SER
AR2-THR
AR2-TR1
AR2-TR2
AR2-TR3
AR2-TY1
AR2-TY2
AR2-TY3
AR2-TY4
AR2-VAL
g
r (Å)
SPICA ver1
SPICA ver2
CHARMM

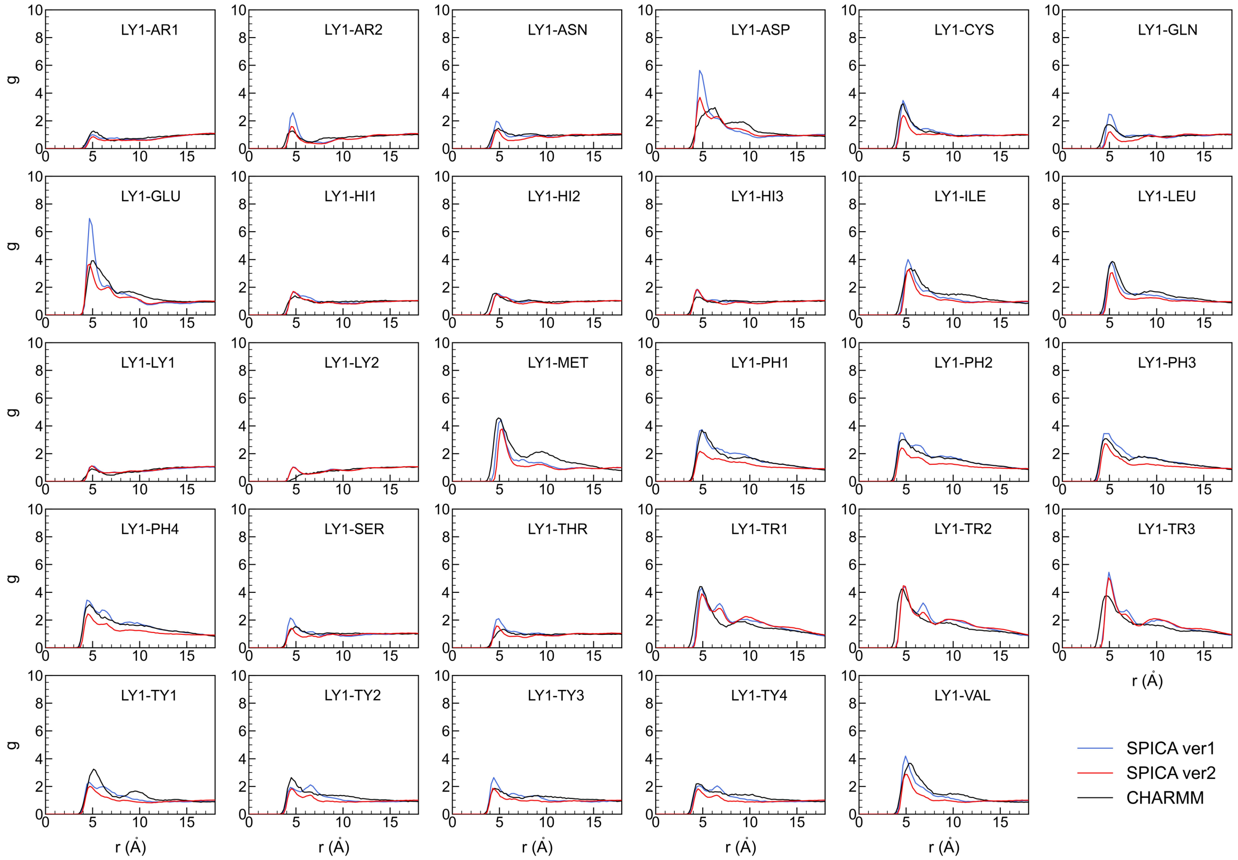
LY1-AR1
LY1-AR2
LY1-ASN
LY1-ASP
LY1-CYS
LY1-GLN
LY1-GLU
LY1-HI1
LY1-HI2
LY1-HI3
LY1-ILE
LY1-LEU
LY1-LY1
LY1-LY2
LY1-MET
LY1-PH1
LY1-PH2
LY1-PH3
LY1-PH4
LY1-SER
LY1-THR
LY1-TR1
LY1-TR2
LY1-TR3
LY1-TY1
LY1-TY2
LY1-TY3
LY1-TY4
LY1-VAL
g
r (Å)
SPICA ver1
SPICA ver2
CHARMM

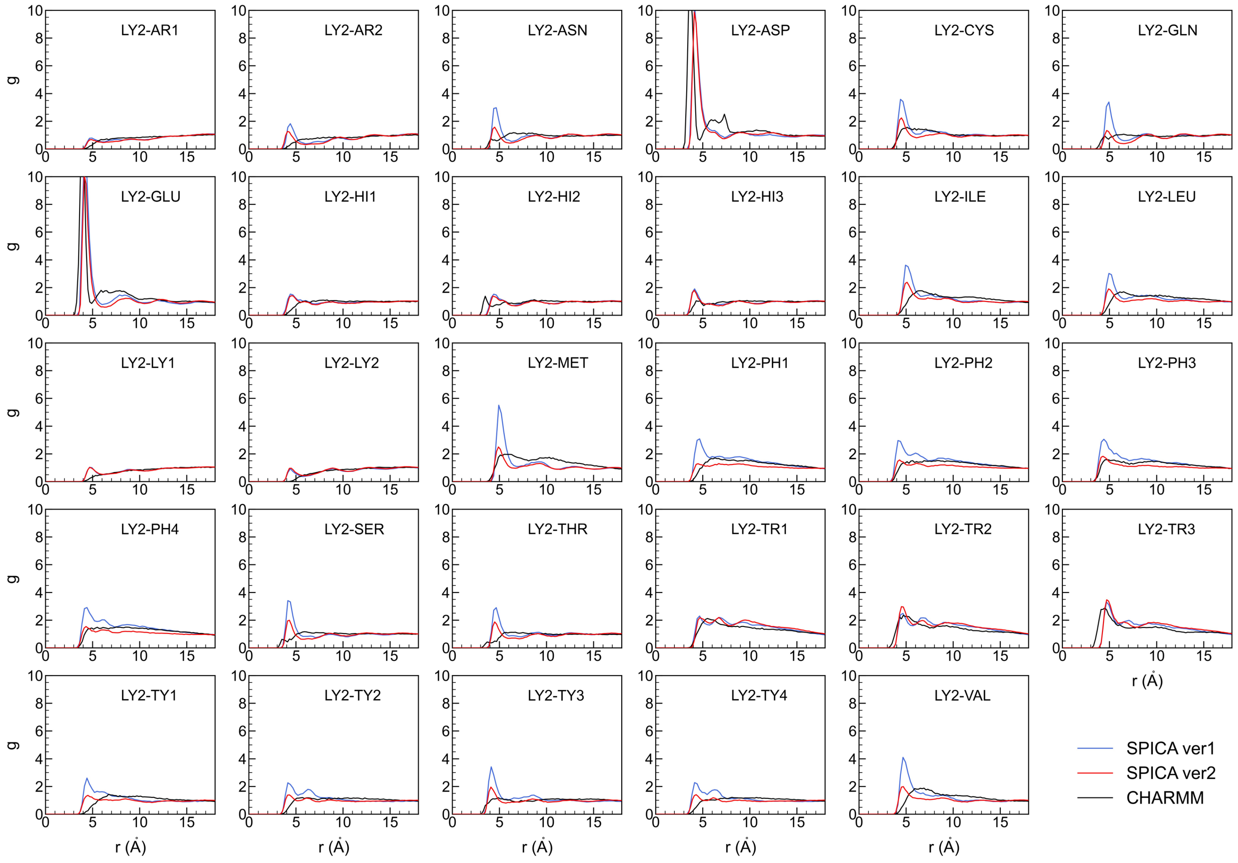
LY2-AR1
LY2-AR2
LY2-ASN
LY2-ASP
LY2-CYS
LY2-GLN
LY2-GLU
LY2-HI1
LY2-HI2
LY2-HI3
LY2-ILE
LY2-LEU
LY2-LY1
LY2-LY2
LY2-MET
LY2-PH1
LY2-PH2
LY2-PH3
LY2-PH4
LY2-SER
LY2-THR
LY2-TR1
LY2-TR2
LY2-TR3
LY2-TY1
LY2-TY2
LY2-TY3
LY2-TY4
LY2-VAL
g
r (Å)
SPICA ver1
SPICA ver2
CHARMM

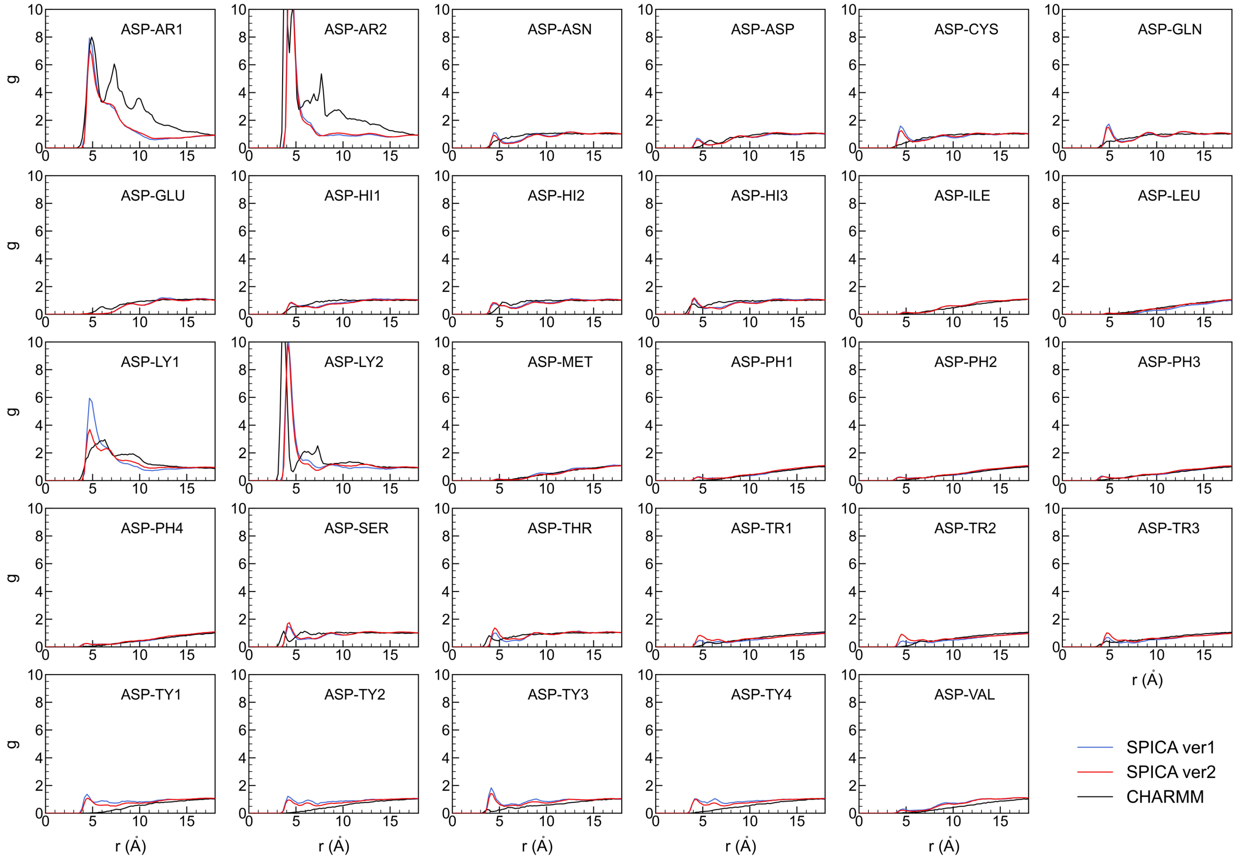
ASP-AR1
ASP-AR2
ASP-ASN
ASP-ASP
ASP-CYS
ASP-GLN
ASP-GLU
ASP-HI1
ASP-HI2
ASP-HI3
ASP-ILE
ASP-LEU
ASP-LY1
ASP-LY2
ASP-MET
ASP-PH1
ASP-PH2
ASP-PH3
ASP-PH4
ASP-SER
ASP-THR
ASP-TR1
ASP-TR2
ASP-TR3
ASP-TY1
ASP-TY2
ASP-TY3
ASP-TY4
ASP-VAL
g
r (Å)
SPICA ver1
SPICA ver2
CHARMM

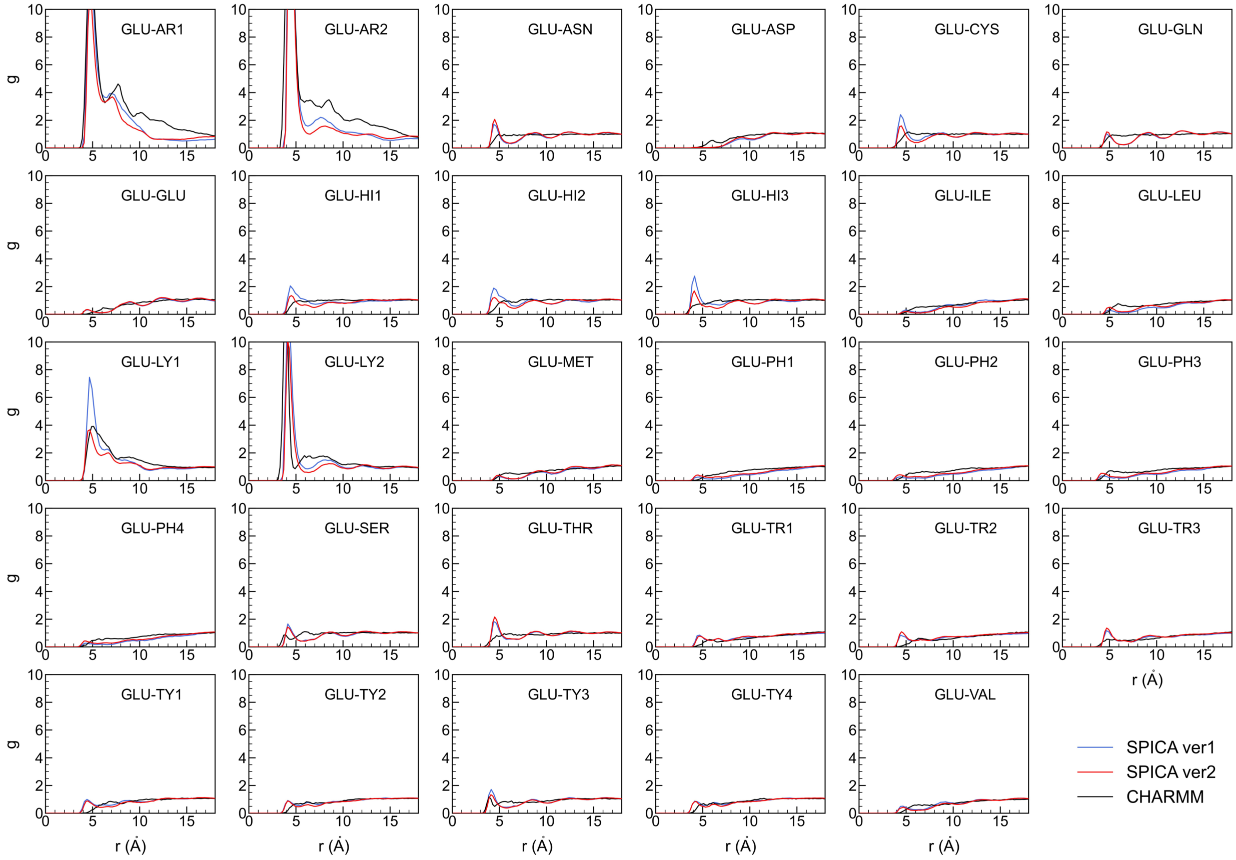
GLU-AR1
GLU-AR2
GLU-ASN
GLU-ASP
GLU-CYS
GLU-GLN
GLU-GLU
GLU-HI1
GLU-HI2
GLU-HI3
GLU-ILE
GLU-LEU
GLU-LY1
GLU-LY2
GLU-MET
GLU-PH1
GLU-PH2
GLU-PH3
GLU-PH4
GLU-SER
GLU-THR
GLU-TR1
GLU-TR2
GLU-TR3
GLU-TY1
GLU-TY2
GLU-TY3
GLU-TY4
GLU-VAL
g
r (Å)
SPICA ver1
SPICA ver2
CHARMM

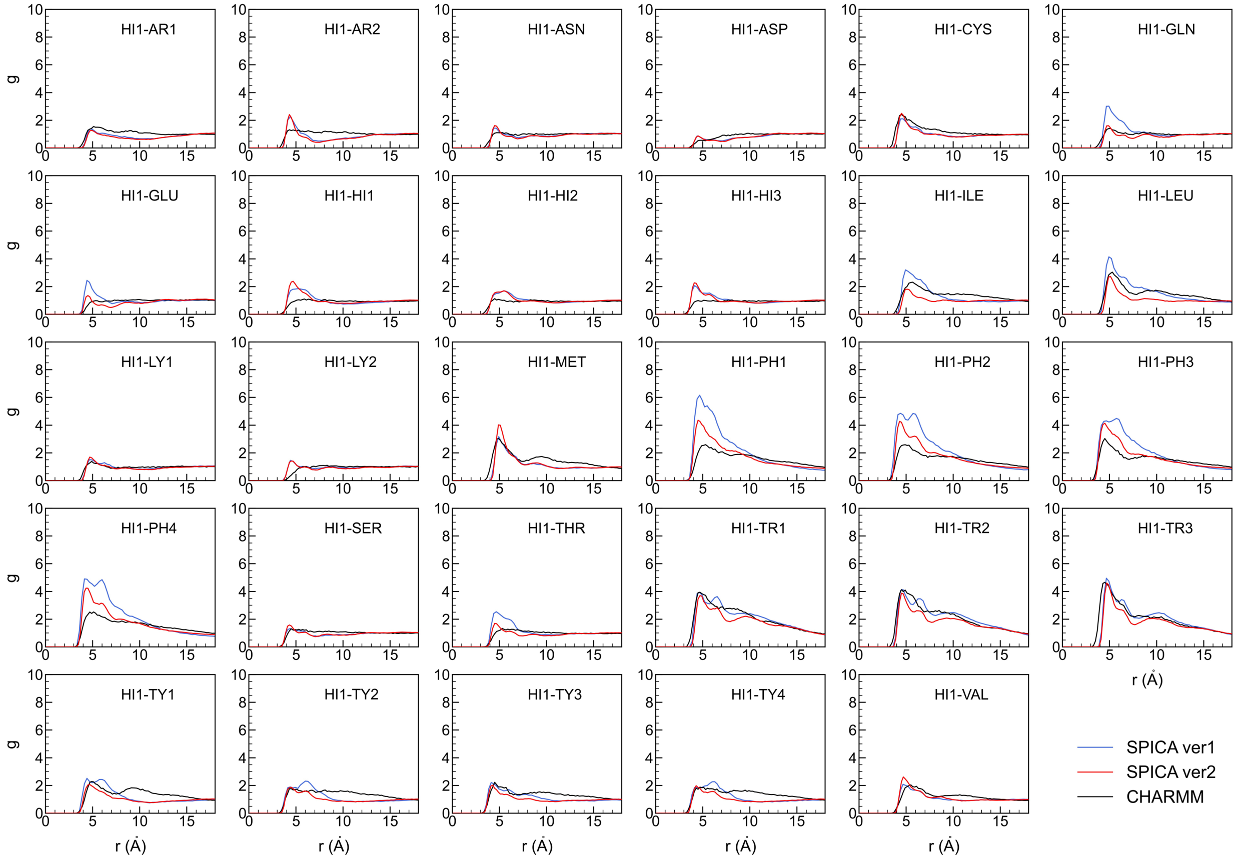
HI1-AR1
HI1-AR2
HI1-ASN
HI1-ASP
HI1-CYS
HI1-GLN
HI1-GLU
HI1-HI1
HI1-HI2
HI1-HI3
HI1-ILE
HI1-LEU
HI1-LY1
HI1-LY2
HI1-MET
HI1-PH1
HI1-PH2
HI1-PH3
HI1-PH4
HI1-SER
HI1-THR
HI1-TR1
HI1-TR2
HI1-TR3
HI1-TY1
HI1-TY2
HI1-TY3
HI1-TY4
HI1-VAL
g
r (Å)
SPICA ver1
SPICA ver2
CHARMM

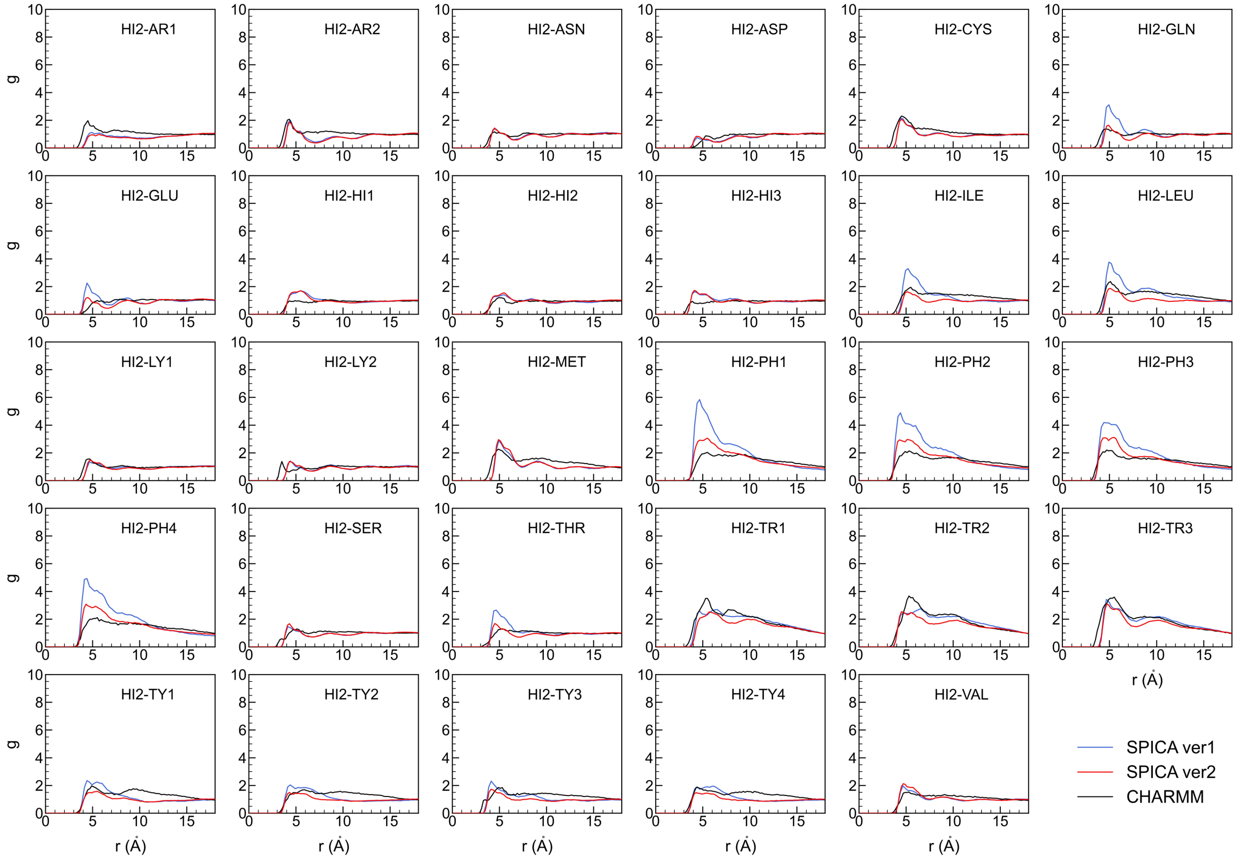
HI2-AR1
HI2-AR2
HI2-ASN
HI2-ASP
HI2-CYS
HI2-GLN
HI2-GLU
HI2-HI1
HI2-HI2
HI2-HI3
HI2-ILE
HI2-LEU
HI2-LY1
HI2-LY2
HI2-MET
HI2-PH1
HI2-PH2
HI2-PH3
HI2-PH4
HI2-SER
HI2-THR
HI2-TR1
HI2-TR2
HI2-TR3
HI2-TY1
HI2-TY2
HI2-TY3
HI2-TY4
HI2-VAL
g
r (Å)
SPICA ver1
SPICA ver2
CHARMM

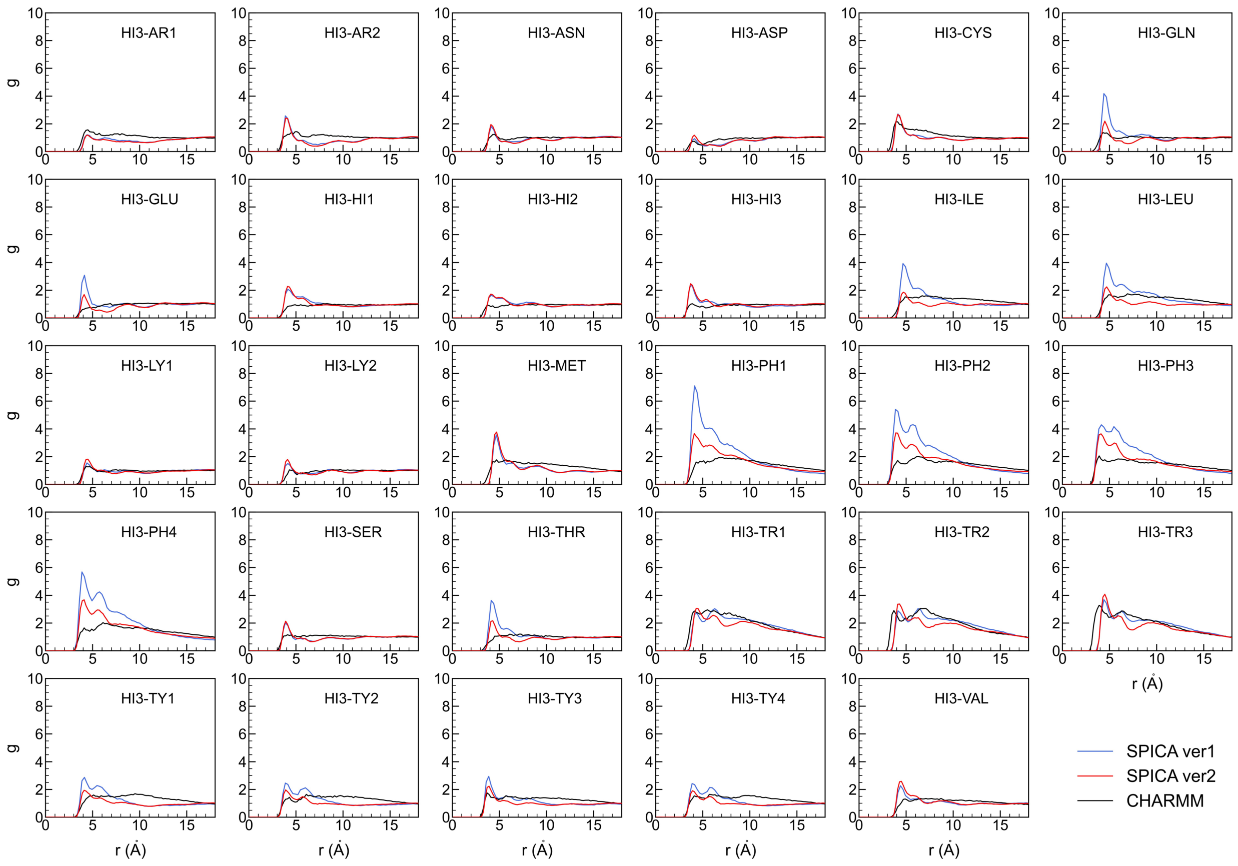


**Figure S2.** Radial distribution functions, *g*, between amino acids side chain segments involving charged one in water.

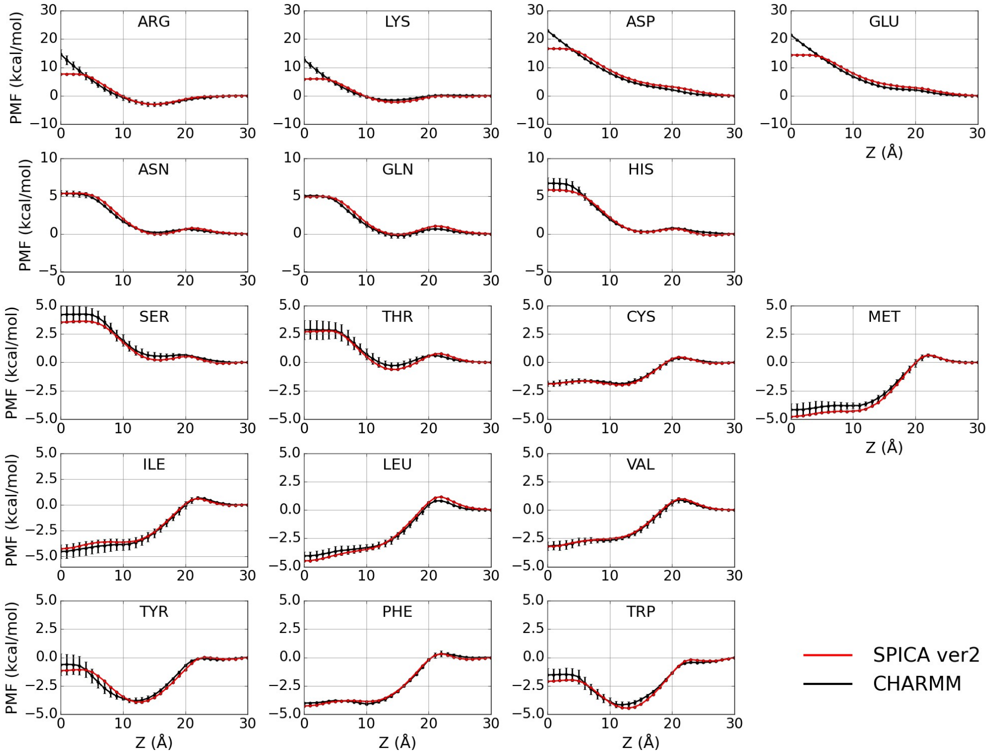


**Figure S3.** The transfer free energy profile of SC analogues across DOPC bilayer.

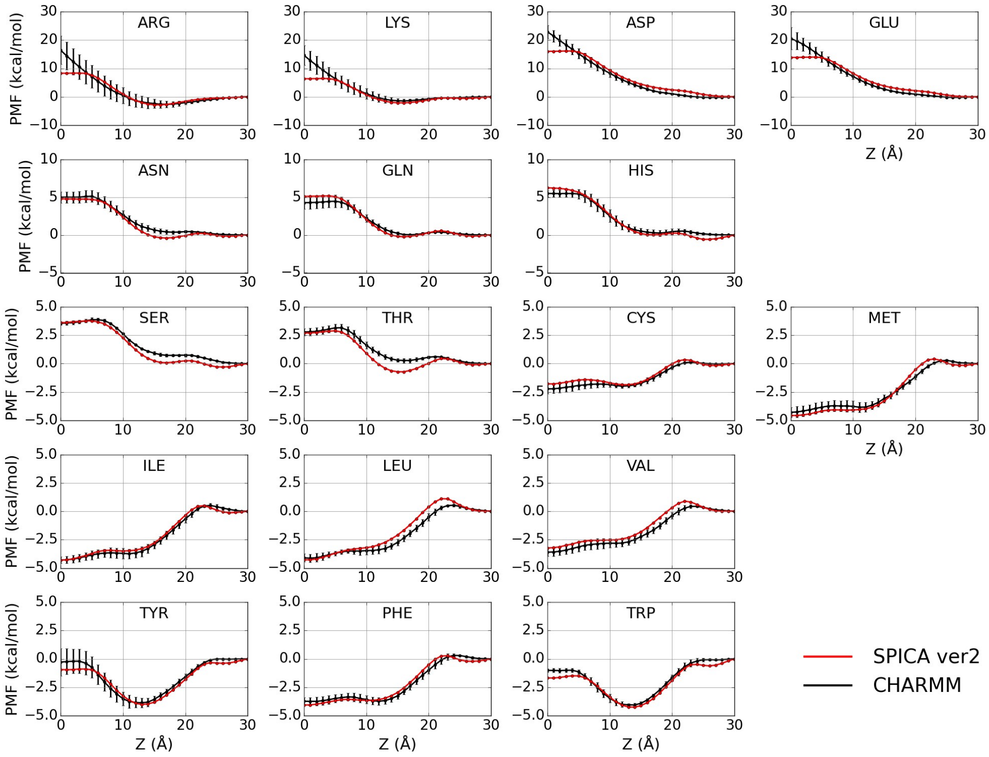


**Figure S4.** The transfer free energy profile of SC analogues across POPE bilayer.

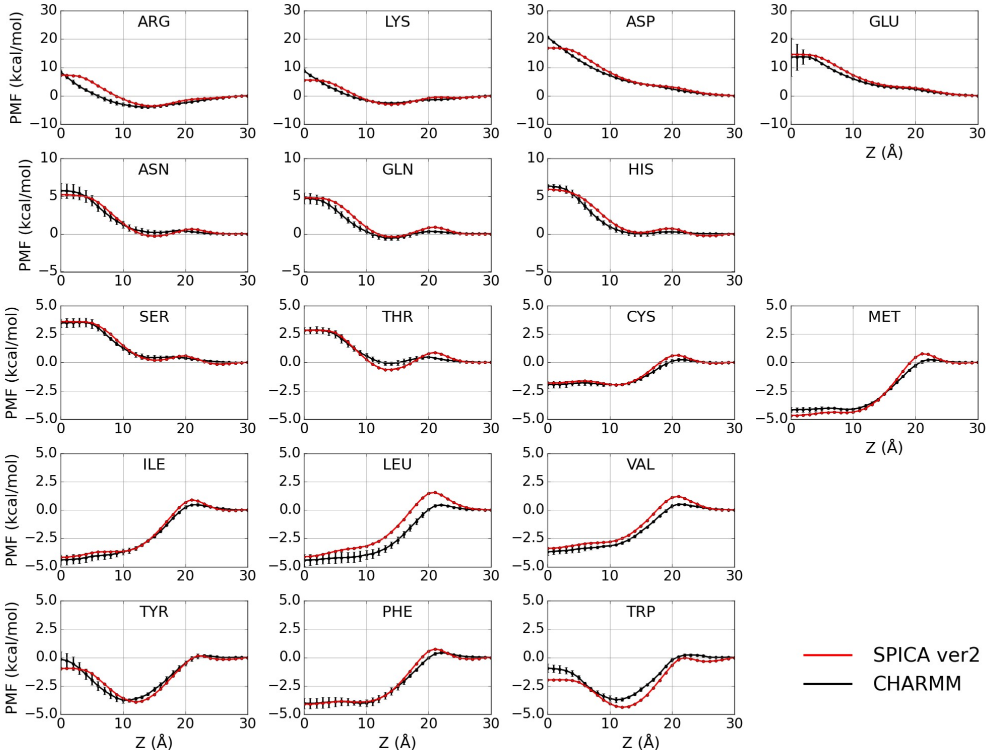


**Figure S5.** The transfer free energy profile of SC analogues across POPG bilayer.

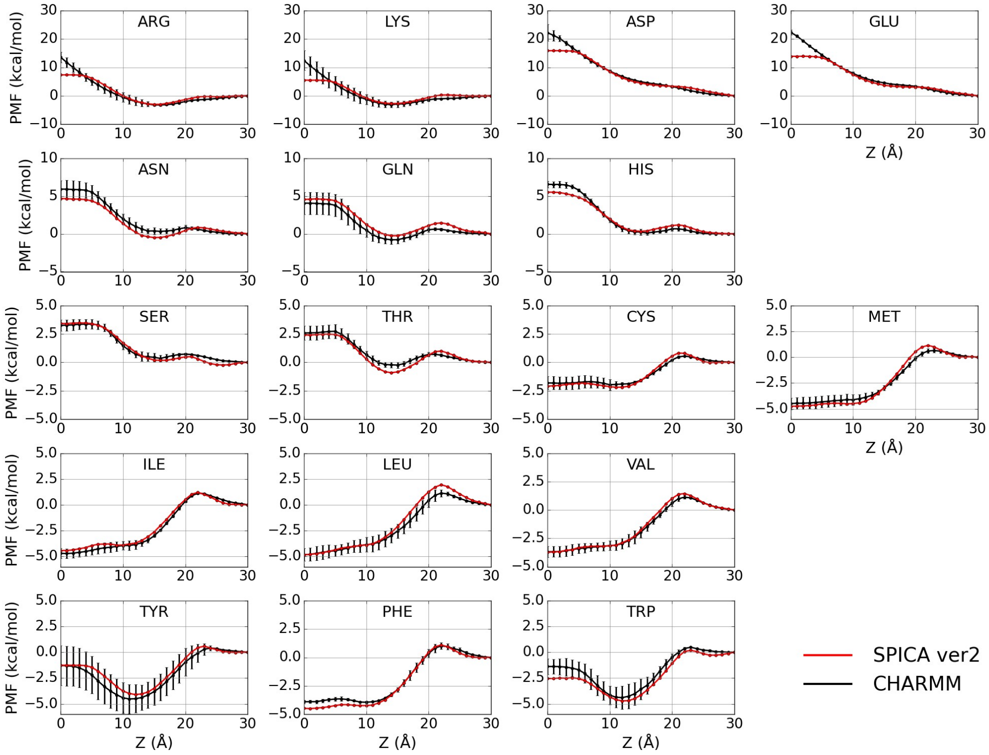


**Figure S6.** The transfer free energy profile of SC analogues across POPI bilayer.

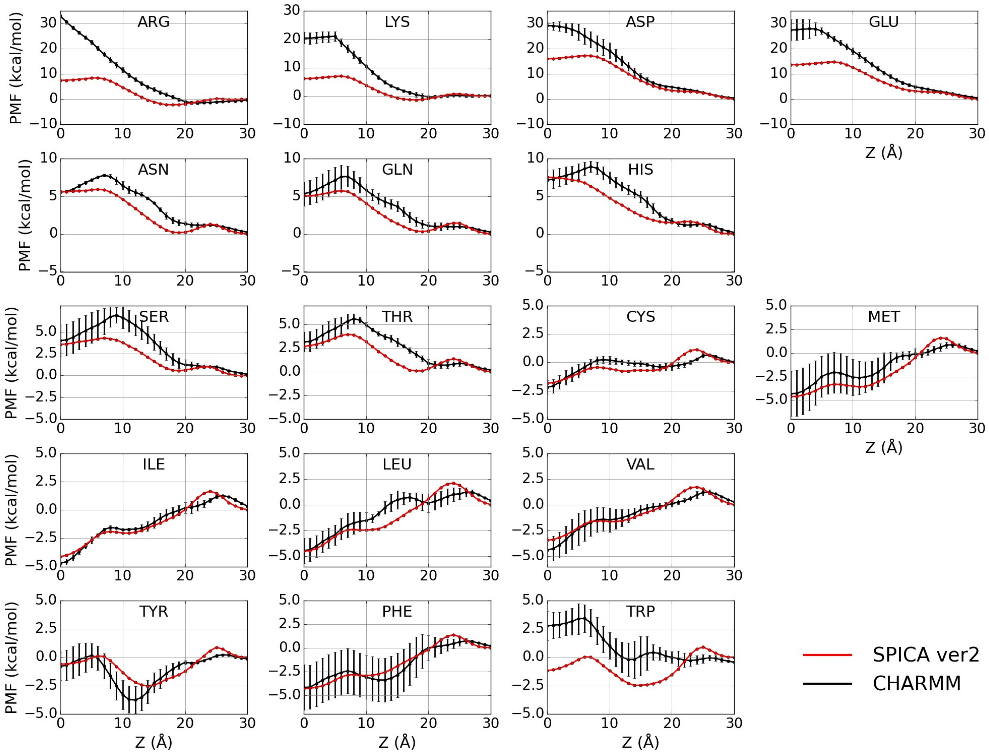


**Figure S7.** The transfer free energy profile of SC analogues across DPPC-CHOL bilayer.

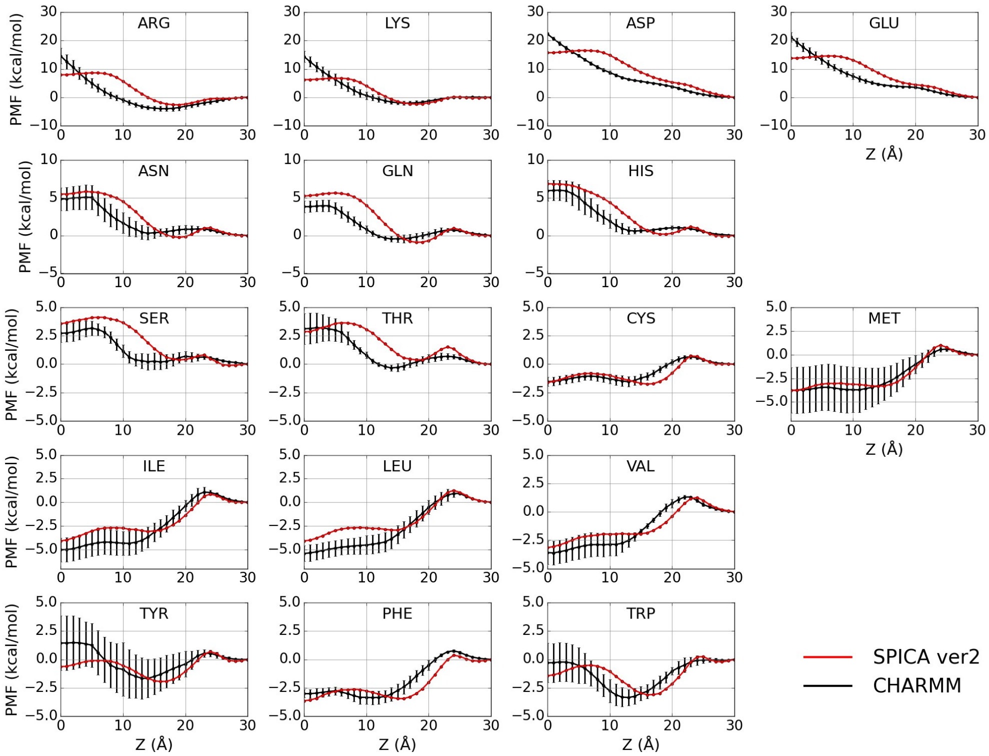


**Figure S8.** The transfer free energy profile of SC analogues across SSM bilayer.

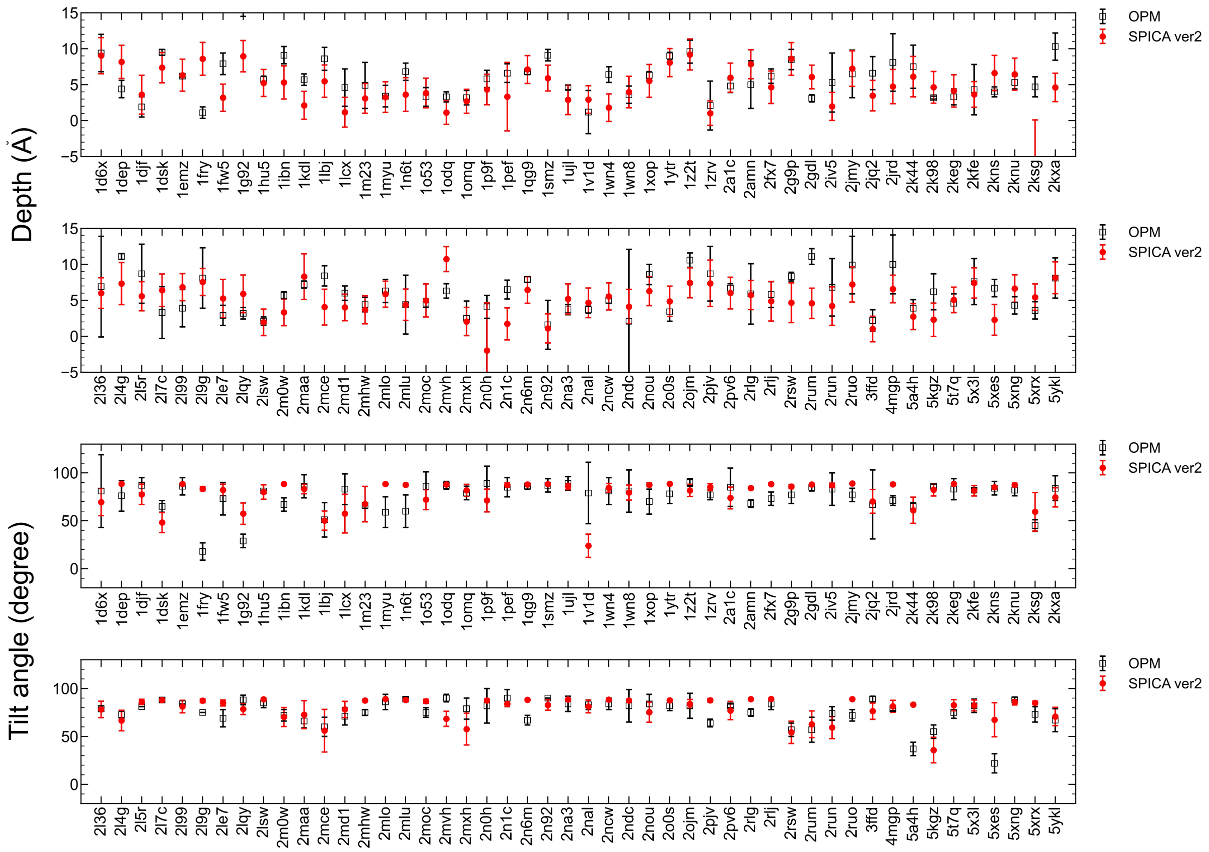


**Figure S9.** The penetration depths and tilt angles of peripheral helices. A comparison of simulated results with SPICA ver2 and OPM database.

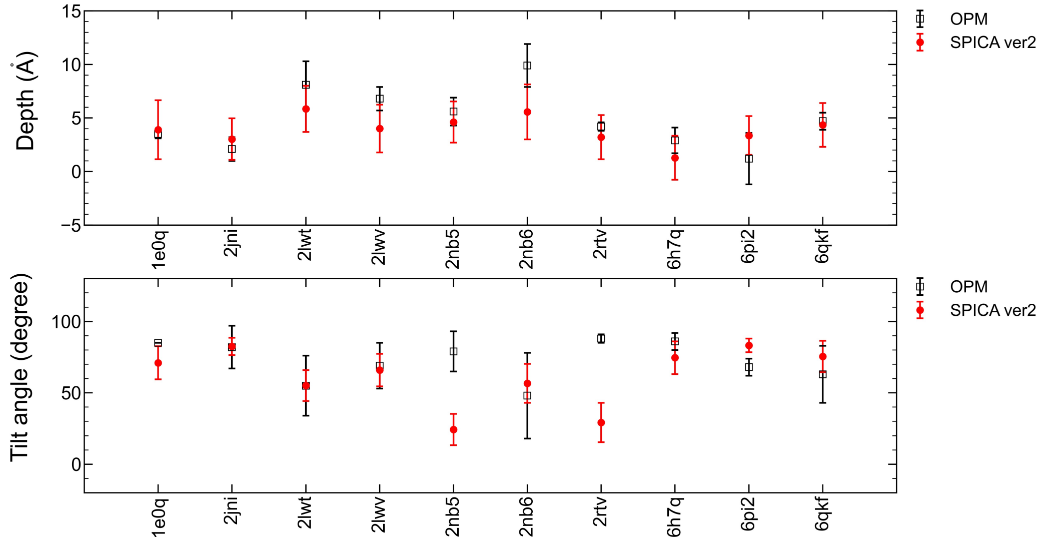


**Figure S10.** The penetration depths and tilt angles of peripheral β-hairpin peptides. A comparison of simulated results with SPICA ver2 and OPM database.
